# Anti-IL1RAP Antibodies for Pan-Inhibition of IL-1 Family Cytokine Signaling in Inflammatory Diseases and Oncology

**DOI:** 10.64898/2026.02.13.705739

**Authors:** Qinghao Liu, Yanxia Wang, Yan Jiang, Yaqi Ru, Qian Zhao, Zhen Zhang, Zhiqiang Li, Qian Shi, Yunlong Cao

## Abstract

The interleukin-1 (IL-1) superfamily of cytokines—comprising the IL-1, IL-33, and IL-36 subfamilies—orchestrates a vast array of innate and adaptive immune responses. While physiological activation of these pathways is essential for host defense and tissue repair, their chronic dysregulation is pathognomonic of a broad spectrum of autoimmune diseases and malignancies. Current therapeutic modalities largely rely on the neutralization of single cytokines, an approach often limited by the functional redundancy and compensatory signaling loops inherent to the IL-1 superfamily. The Interleukin-1 Receptor Accessory Protein (IL1RAP) serves as an obligatory co-receptor for IL-1, IL-33, and IL-36 subfamilies, functioning as a molecular bottleneck for pro-inflammatory signaling. Herein, we report the development and comprehensive characterization of two humanized monoclonal antibodies, DXP-006 and DXP-106, which target a unique epitope on domain 2 of IL1RAP. Cryo-electron microscopy studies at 3.55 Å resolution reveal that these antibodies lock the critical c2d2 loop of IL1RAP, sterically occluding its recruitment to IL-1R1, ST2 and IL-36R binary complexes and potently blockade IL-1, IL33, and IL-36 signaling. DXP-006, engineered with a half-life-extended and effector-silenced Fc, exerts profound efficacy in murine models of psoriasis, superior to the clinical benchmark bimekizumab. Conversely, DXP-106, engineered with a defucosylated Fc to enhance antibody-dependent cellular cytotoxicity, exhibits potent antitumor activity in non-small cell lung cancer xenografts. These findings position DXP-006 and DXP-106 as ideal therapeutics for IL-1 family-driven autoimmunity and IL1RAP-positive cancers.

**One Sentence Summary:** Anti-IL1RAP antibodies block multiple IL-1 family cytokines simultaneously, showing superior efficacy in inflammation and cancer models.

## INTRODUCTION

The vertebrate immune system relies on a highly sophisticated, evolutionarily ancient network of cytokines to initiate inflammation in response to injury or infection. Among these, the Interleukin-1 (IL-1) superfamily represents one of the most potent signaling networks (1,2), orchestrating the critical transition from innate to adaptive immunity, driving the clonal expansion of T-helper subsets, and regulating the activation of epithelial and endothelial barriers. While the physiological activation of these pathways is essential for host defense, their chronic dysregulation drives a broad spectrum of human pathologies, ranging from autoinflammatory syndromes and chronic autoimmune diseases to the progression of solid tumors where inflammation fuels angiogenesis and immune evasion.

The complexity of this superfamily presents both a therapeutic opportunity and a formidable challenge due to the significant functional redundancy inherent to its architecture. The superfamily is broadly categorized into subfamilies based on receptor utilization. The IL-1 subfamily, including IL-1α and IL-1β, binds to the type 1 IL-1 receptor (IL-1R1) and is central to rheumatoid arthritis and gout (2,3). The IL-33 subfamily functions as an alarmin released upon epithelial stress, binding to the ST2 receptor to drive Type 2 immune responses implicated in asthma and fibrosis (4). The IL-36 subfamily comprises three agonists (IL-36α, IL-36β, IL-36γ) that bind to the IL-36 receptor, maintaining skin homeostasis but driving severe pathology in generalized pustular psoriasis (GPP) and inflammatory bowel disease (IBD) (5).

Importantly, these cytokines often act synergistically, exacerbating disease severity when multiple members are dysregulated. In psoriasis, IL1β and IL36γ cooperatively enhance keratinocyte activation (6), while in RA, IL33 and IL1β jointly promote fibroblast proliferation and synovitis (3). The current standard of care for many autoimmune diseases involves biologic agents that target individual cytokines. Canakinumab (anti-IL-1β) and anakinra (IL-1Ra) are effective in specific autoinflammatory diseases like Cryopyrin-Associated Periodic Syndromes (CAPS) but have shown limited or mixed efficacy in broader, heterogeneous diseases like severe psoriasis, systemic sclerosis (SSc), or chronic obstructive pulmonary disease (COPD) (7). Similarly, specific inhibition of IL-36R (spesolimab) is highly effective for acute flares of GPP but may not fully resolve chronic plaque psoriasis where IL-1 and IL-17 axes are dominant (8). Thus, simultaneous blockade of multiple IL-1 family cytokines may offer superior therapeutic efficacy compared to single-target inhibition.

Despite utilizing distinct primary receptors (IL-1R1, ST2, and IL-36R) for ligand capture, all three subfamilies require the subsequent recruitment of a shared, obligate co-receptor: the Interleukin-1 Receptor Accessory Protein (IL1RAP) (also known as IL-1R3 or IL-1RAcP) (9). IL1α and IL1β signal through IL1R1 and IL1RAP (10), while IL33 utilizes ST2 (IL-1RL1) and IL1RAP (11). IL-36 cytokines (IL-36α, β, γ) bind IL-36R and also require IL1RAP for downstream NF-κB and MAPK activation (12). Because IL1RAP is the common denominator for six distinct pro-inflammatory cytokines, it represents a high-value therapeutic target. Blocking IL1RAP theoretically offers “pan-IL-1 family inhibition,” a strategy that could overcome the limitations of single-cytokine blockade observed in the clinic (13).

Beyond autoimmune diseases, IL1RAP is increasingly recognized as a key player in tumor progression, particularly in solid malignancies (14,15). The IL-1 family cytokines contribute to a pro-tumorigenic microenvironment by promoting angiogenesis, immune evasion, and metastasis (16). IL-1Ra and anti-IL-1 mAbs have been shown to inhibit primary tumor growth and metastasis (17). The potential for IL-1 blockade to synergize with checkpoint inhibitors has been studied in various models (18). IL1RAP is overexpressed in several cancers, including hematological cancers and solid cancers (15), where it facilitates tumor cell survival and resistance to therapy (19,20,21), making it a potential tumor-associated antigen (TAA). Given these roles, IL1RAP blockade represents a dual therapeutic strategy—modulation of cancer-associated inflammation and targeted therapy.

The validation of IL1RAP as a therapeutic target has catalyzed the development of several clinical-stage antibodies, including Nadunolimab and CAN10. Nadunolimab is a humanized IgG1 antibody currently in Phase II trials for pancreatic ductal adenocarcinoma (PDAC) and NSCLC (22). Its primary mechanism of action involves blocking IL-1α and IL-1β signaling and inducing Antibody-Dependent Cellular Cytotoxicity (ADCC) against IL1RAP-expressing tumor cells. However, its inhibitory spectrum is largely limited to IL-1 and IL-36, with minimal activity against IL33. This leaves the potential for IL33 mediated escape mechanisms in the TME. CAN10, a broad-spectrum anti-IL-1RAP antibody, blocks all six IL-1RAP-dependent cytokines (IL1α, IL1β, IL33, IL36α/β/γ) (23,24). While broader in spectrum than Nadunolimab, its development focuses on autoimmune and fibrotic indications (e.g., systemic sclerosis, myocarditis). The affinity and epitope engagement of CAN10 have set a benchmark, yet comparative structural analysis suggests room for optimization in binding kinetics and epitope “locking” efficiency to achieve maximal potency.

These findings underscored the need for improved IL1RAP inhibitors with higher affinity and broader neutralizing capacity. To address the limitations of current IL1RAP-targeted therapies, we developed two novel high-affinity antibodies, DXP-006 and DXP-106, which “lock” the c2d2 loop of IL1RAP to potent blockade of IL-1, IL33, and IL-36 signaling. Our engineering strategy was bifurcated to meet the distinct biological requirements of autoimmunity versus oncology. DXP-006 was engineered with Fc mutations to silence effector functions and extend serum half-life, making it suitable for chronic autoimmunity where target neutralization is the goal. Conversely, DXP-106 was engineered with a defucosylated Fc to dramatically enhance antibody-dependent cellular cytotoxicity (ADCC), aiming to physically eliminate IL1RAP-expressing tumor cells in oncology settings. Preclinical *in vitro* and *in vivo* experiments position DXP-006 and DXP-106 as a potentially best-in-class candidate for oncological indications, offering broader and more durable cytokine inhibition and more efficient than current alternatives.

## RESULTS

### High-Affinity Antibodies Binding to IL1RAP

The extracellular domain (ECD) of IL1RAP consists of three Immunoglobulin-like (Ig-like) domains: D1, D2, and D3. Structural biology studies of the IL-1β signaling complex (PDB: 4DEP) have established that Domain 2 (D2) is the critical interface for recruitment to the IL-1R1 binary complex. Within D2, a region termed the c2d2 loop (connecting the c2 and d2 -strands of Domain 2) functions as a hydrophobic latch that engages the binary complex (Figure 1A and S1A-1C). To generate antibodies with potent pan-inhibitory activity, we prioritized binders that specifically target this functional D2 epitope. We screened antibodies derived from rabbits and identified one parental antibody that binds to domain 2 (Figure S1D and S1E). ELISA assays showed that the parental antibody bound to neither domain 1 nor domain 3 of IL1RAP but did bind to the full ECD and to the domain 1+domain 2 construct, confirming its domain specificity.

**Fig. 1.**
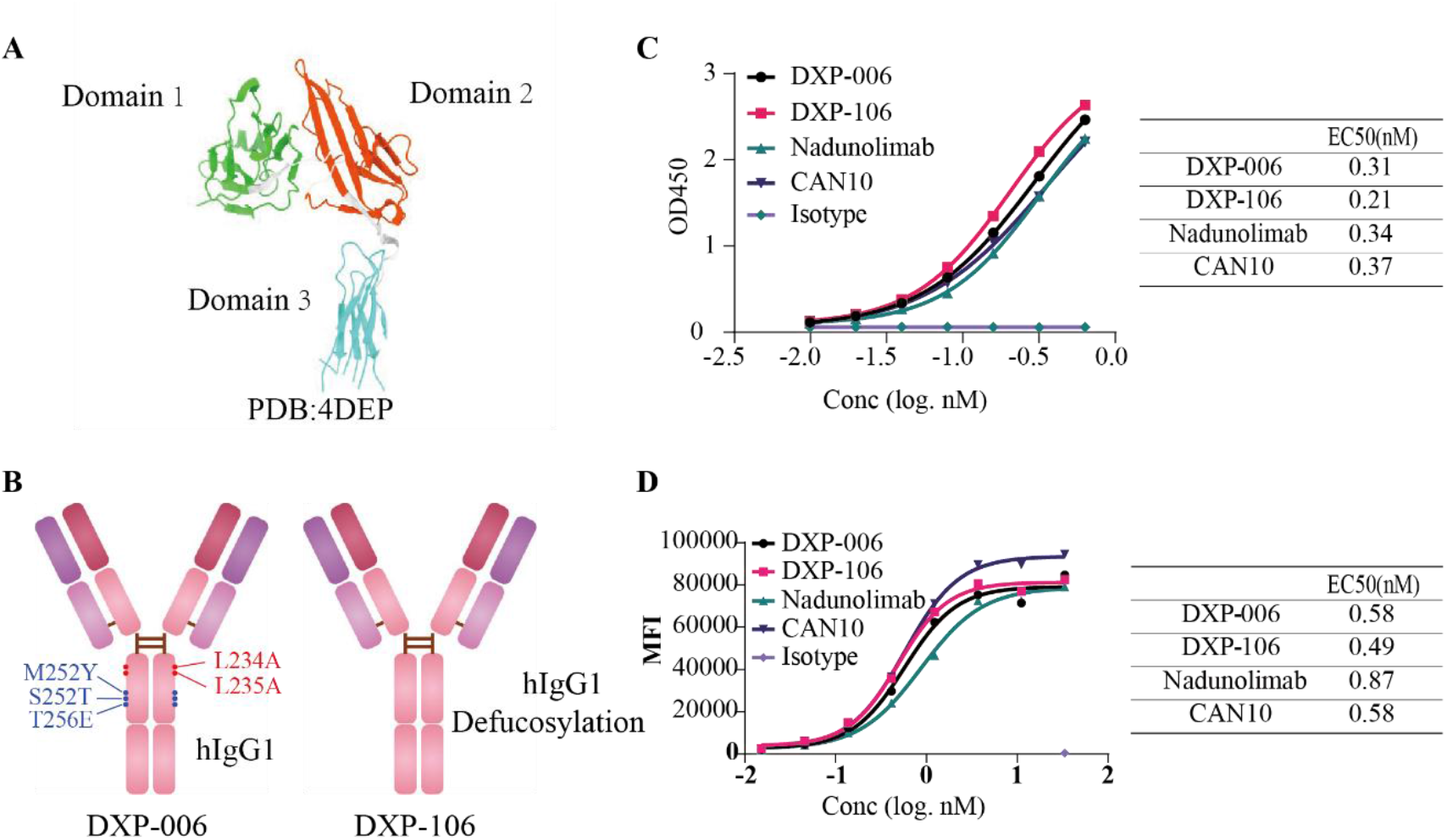
DXP-006 and DXP-106 bind with high affinity to IL1RAP. (A) Schematic diagram of IL1RAP ECD structure. (B) Schematic diagram of DXP-006 and DXP-106 antibody structures. (C) ELISA binding curves showing dose-dependent binding of DXP-006, DXP-106 to full-length IL1RAP. (D) Flow cytometry analysis of DXP-006, DXP-106 and benchmark antibodies binding to HEK293T cells overexpressing human IL1RAP.

To minimize immunogenicity, we humanized the parental antibody, generating two variants: DXP-006 and DXP-106 (Figure 1B). For chronic autoimmune conditions, the therapeutic goal is pure receptor blockade without depleting host immune cells. Consequently, the Fc region of DXP-006 (human IgG1) was engineered with L234A and L235A mutations (LALA) to silence effector functions, preventing ADCC and CDC. Additionally, DXP-006 incorporates M252Y/S254T/T256E (YTE) mutations to increase affinity for the neonatal Fc receptor (FcRn) at acidic pH, promoting recycling and extending serum half-life. Conversely, for oncology, where IL1RAP acts as a tumor-associated antigen, DXP-106 is purposefully designed with a defucosylated Fc architecture, dramatically increasing its affinity for FcγRIIIa (CD16a) on Natural Killer (NK) cells to maximize ADCC potency.

Bio-Layer Interferometry (BLI) confirmed that both DXP-006 and DXP-106 bind human IL1RAP with sub-nanomolar affinity (Figure S2A; Table S1). In ELISA assays, both DXP-006 and DXP-106 also demonstrated high affinity for IL1RAP, with EC50 values 1.2-fold and 1.6-fold lower than that of CAN10 and Nadunolimab (Figure 1C). Furthermore, we utilized the IL1RAP-overexpressing 293T cell line hIL1RAP/293T to evaluate the affinity to the cell surface IL1RAP. DXP-006 and DXP-106 both exhibit potent binding to membrane-bound IL1RAP, with EC50 values superior or similar to those of the control antibodies CAN10 and Nadunolimab (Figure 1D).

Given the high sequence homology between human and cynomolgus IL1RAP ECDs, we assessed the cynomolgus cross-reactivity of DXP-006 and DXP-106. BLI analysis revealed that both DXP-006 and DXP-106 retain high affinity for cynomolgus IL1RAP with KD values at the nanomolar level (Figure S2B; Table 1). The high affinity for cynomolgus monkey IL1RAP validates the use of this species for toxicology studies. Collectively, these data establish DXP-006 and DXP-106 as high-affinity antibodies that selectively target domain 2 of human and cynomolgus IL1RAP.

**Table 1.**
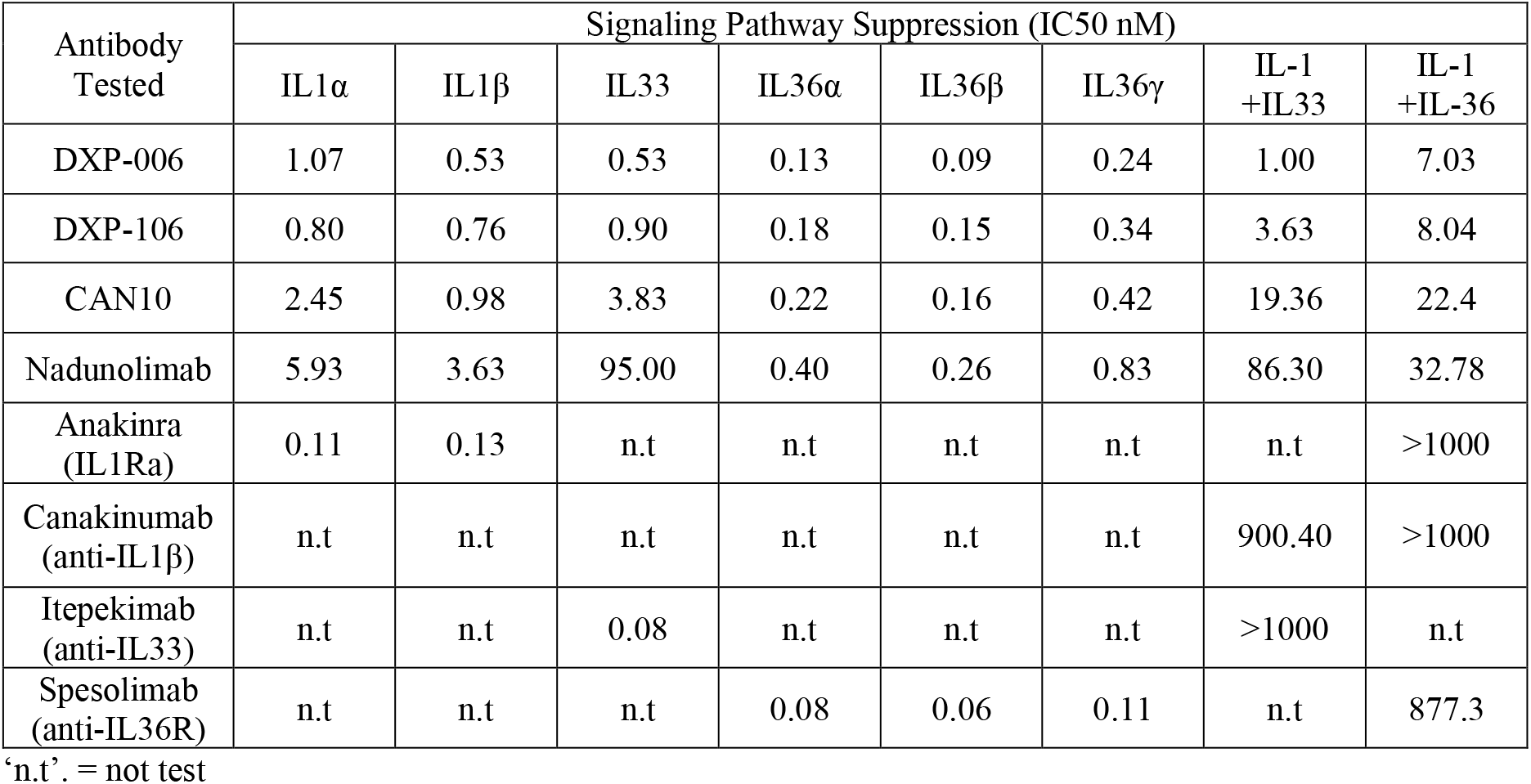
Summary of inhibitory potency across cytokine assays.

| Antibody Tested | Signaling Pathway Suppression (IC50 nM) |  |  |  |  |  |  |  |
| --- | --- | --- | --- | --- | --- | --- | --- | --- |
| | IL1α | IL1β | IL33 | IL36α | IL36β | IL36γ | IL-1 +IL33 | IL-1 +IL-36 |
| DXP-006 | 1.07 | 0.53 | 0.53 | 0.13 | 0.09 | 0.24 | 1.00 | 7.03 |
| DXP-106 | 0.80 | 0.76 | 0.90 | 0.18 | 0.15 | 0.34 | 3.63 | 8.04 |
| CAN10 | 2.45 | 0.98 | 3.83 | 0.22 | 0.16 | 0.42 | 19.36 | 22.4 |
| Nadunolimab | 5.93 | 3.63 | 95.00 | 0.40 | 0.26 | 0.83 | 86.30 | 32.78 |
| Anakinra (IL1Ra) | 0.11 | 0.13 | n.t | n.t | n.t | n.t | n.t | >1000 |
| Canakinumab (anti-IL1β) | n.t | n.t | n.t | n.t | n.t | n.t | 900.40 | >1000 |
| Itepekimab (anti-IL33) | n.t | n.t | 0.08 | n.t | n.t | n.t | >1000 | n.t |
| Spesolimab (anti-IL36R) | n.t | n.t | n.t | 0.08 | 0.06 | 0.11 | n.t | 877.3 |

‘n.t’. = not test

### Simultaneous Suppression of IL-1 Family Cytokines

IL1RAP serves as a co-receptor for IL-1 (IL1α, IL1β), IL33, and IL-36 (IL36α, IL36β, IL36γ) cytokines, facilitating downstream signaling for all six cytokines. We therefore evaluated the potency of DXP-006 and DXP-106 in functional cellular assays, benchmarking them against clinical standards. HEK293/AP1-NFkB-LUC cells expressing IL1R1 and IL1RAP were utilized to evaluate the inhibitory effects of DXP-006 and DXP-106 on IL1α and IL1β induced activation. IL1R1 antagonist Anakinra can effectively inhibit both IL1α and IL1β mediated activation of NF-κB/AP-1. DXP-006 and DXP-106 potently inhibited IL1α signaling, demonstrating 2.3-fold and 7.4-fold lower IC50 values compared to the control antibodies CAN10 and Nadunolimab under identical experimental conditions (Figure 2A, Table 1). Similarly, in the IL1β pathway, IC50 values for DXP-006 and DXP-106 were 1.8-fold and 4.8-fold lower than the controls (Figure 2B, Table 1).

**Fig. 2.**
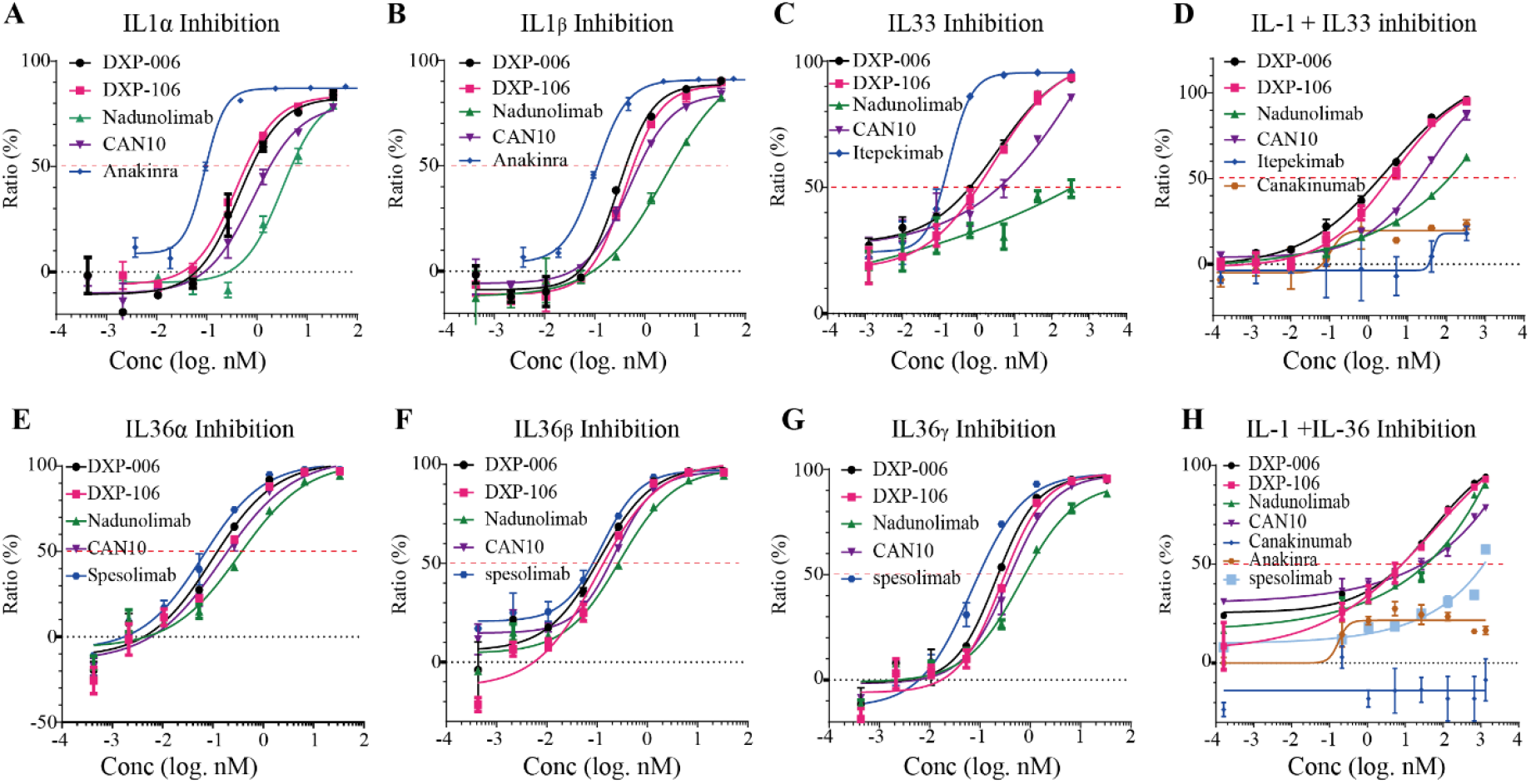
DXP-006 and DXP-106 potently inhibit signaling across all three IL-1 subfamilies. (A-B) Inhibition of IL1α (A) and IL1β (B) induced signaling by indicated antibodies in HEK-Blue IL-1 reporter cells. (C) Inhibition of IL33 induced signaling by indicated antibodies in HUVEC cells. (D) Inhibition of IL1α, IL1β, and IL33 induced signaling by indicated antibodies in HUVEC cells. (E-G) Inhibition of IL36α (E), IL36β (F) and IL36γ (G) induced signaling by indicated antibodies in A431 cells. (H) Inhibition of IL1α, IL1β, IL36α, IL36β, and IL36γ induced signaling by indicated antibodies in A431 cells.

Next, we assessed the inhibitory effects of DXP-006 and DXP-106 on IL33 signaling in human umbilical vein endothelial cells (HUVEC). The Th2-polarizing cytokine IL33 triggers IL-8 secretion from HUVEC (25), which can be suppressed by the IL33 monoclonal antibody Itepekimab. DXP-006 and DXP-106 effectively inhibit IL33-induced IL8 secretion in HUVEC and demonstrate superior inhibitory efficacy compared to CAN10 and Nadunolimab (Figure 2C, Table 1). To mimic the complex inflammatory milieu of diseases like psoriasis, we stimulated cells with cytokine “cocktails.” When HUVEC were simultaneously stimulated with IL1α, IL1β, and IL33, a higher level of IL8 secretion was observed. This secretion was only partially inhibited by the IL33 monoclonal antibody Itepekimab or the IL1β monoclonal antibody Canakinumab. By blocking IL1RAP signaling, DXP-006 and DXP-106 potently inhibited IL8 release induced by IL1α, IL1β, and IL33, demonstrating superior efficacy to Nadunolimab and CAN10 (Figure 2D, Table 1).

The inhibitory activity against IL-36 cytokines was assessed in A431 cells, which endogenously express IL36R, IL1RAP, and IL1R1. (26). IL36α, IL36β, and IL36γ-induced activation of A431 cells can all be inhibited by the IL36R antagonist antibody Spesolimab. In the context of IL36α signaling, DXP-006 demonstrates 1.7-fold superior inhibitory activity in IC50 values compared to CAN10, while DXP-106 shows 2.2-fold greater potency in IC50 values relative to Nadunolimab (Figure 2E, Table 1). For IL36β signaling, DXP-006 and DXP-106 demonstrate 1.8-fold and 1.7-fold superior inhibitory activity (IC50 values) compared to CAN10 and Nadunolimab, respectively (Figure 2F, Table 1). In the case of IL36γ signaling, DXP-006 and DXP-106 also demonstrate potent inhibitory activity, with IC50 values that are 1.8-fold and 2.4-fold superior to those of CAN10 and Nadunolimab (Figure 2G, Table 1). The inhibition of the synergistic biological activity between IL-1 and IL-36 by DXP-006 and DXP-106 was also assessed in A431 cells. When A431 cells were simultaneously stimulated with the five cytokines, it was found that neither the IL1β mAb Canakinumab, the IL1R1 antagonist Anakinra, nor the IL36R antagonist antibody Spesolimab could effectively suppress IL8 secretion. By blocking IL1RAP signaling, DXP-006 and DXP-106 potently suppress IL8 release induced by IL1α, IL1β, IL36α, IL36β, and IL36γ, demonstrating superior activity to Nadunolimab and CAN10 (Figure 2H, Table 1).

In summary, DXP-006 and DXP-106 consistently demonstrate superior inhibitory potency over the control antibodies Nadunolimab and CAN10 against both individual IL1RAP-associated cytokine signaling and combined cytokines signaling. These results underscore the clinical value of pan-inhibition: removing the common denominator IL1RAP collapses the synergistic network that single-cytokine inhibitors fail to address.

### Locking c2d2 Loop to Block IL-1 Family Signaling

To elucidate the molecular mechanism underlying broad inhibition, we determined the cryo-electron microscopy (cryo-EM) structure of the DXP-006/IL1RAP complex (Fig. S3). The resulting electron density map exhibited well-defined features at the antibody–receptor interface, facilitating high-resolution structural analysis (Fig. 3A). Structural analysis revealed two primary interaction interfaces between DXP-006 and IL1RAP. The c2d2 loop and the 152–157 loop of IL1RAP constitute the core contact regions. The heavy chain of DXP-006 engages residues G154, E152, N186, N188, N189, and L200 of IL1RAP, while the light chain interacts with R157, S198, and E193 (Fig. 3B). Notably, the epitope overlaps with that reported for CAN10, which also targets the c2d2 loop and shares key contact residues (N186, N188) (14). Therefore, DXP-006 shares a highly similar IL1RAP binding epitope with CAN10 (Fig. 3C).

**Fig. 3.**
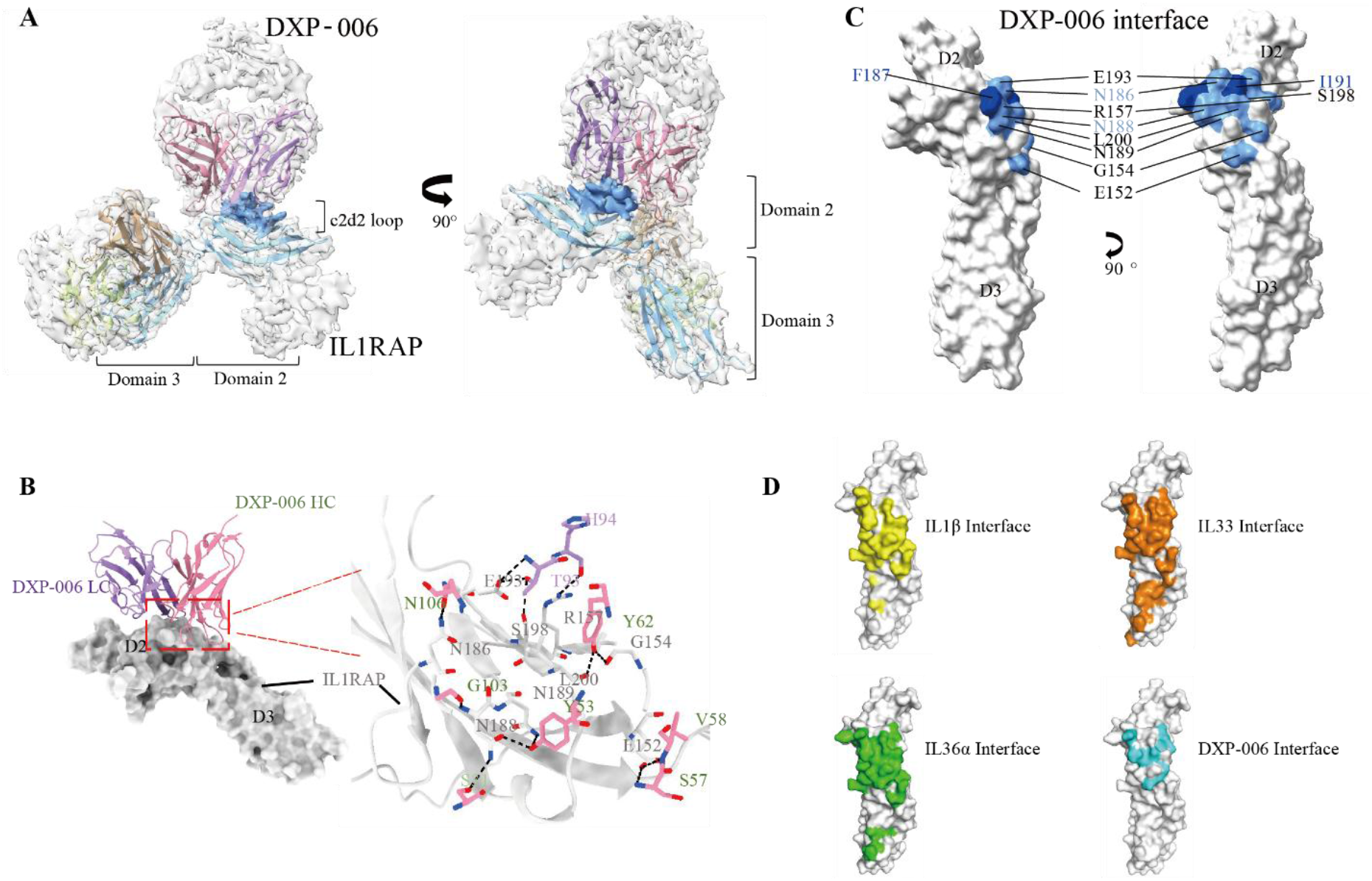
Cryo-EM structure of the DXP-006/IL1RAP complex. (A) Cryo-EM map of the complex that contains DXP-006 (purple and pink), IL1RAP (cyan; domain 2 and domain 3) and density map (grey), c2d2 loop (blue density map). (B) Interfaces between DXP-006 and IL1RAP. (C) Crystal structure of IL1RAP with the DXP-006 Interface (cyan) and CAN10 interface (blue). (D) Crystal structure of IL1RAP domain 2 and domain 3 (PDB:4DEP), colored with IL-1 (yellow), IL33 (orange), IL-36 (green) and DXP-006 (cyan) interface.

Comparative analysis with the IL1β-IL1R1 signaling complex (PDB: 4DEP) shows substantial overlap in binding residues: both DXP-006 and IL1β simultaneously contact E152, N186, and N188 on IL1RAP, while DXP-006 and IL1R1 both engage G154 and N188. The exact structures of IL33 and IL-36 cytokine complexes are not yet available. However, based on similarities, the c2d2 loop is also a key structural component through which IL1RAP participates in complex formation. Therefore, by binding to the c2d2 loop, DXP-006 can effectively block signaling of IL-1, IL33, and IL-36 (Fig. 3D).

### Potent Efficacy of DXP-006 in Inflammatory Disease Models

We evaluated the therapeutic potential of IL1RAP blockade in murine models of inflammatory disease and cancer. Two complementary models were utilized to evaluate DXP-006 in psoriasis-like skin inflammation. C57BL/6 mice were shaved on the dorsal skin two days in advance, and 62.5 mg of imiquimod (IMQ) cream was applied daily to the dorsal skin for eight consecutive days to establish the psoriasis model. LAD191, which is another IL1RAP antagonist monoclonal antibody under clinical evaluation, exhibits stronger inhibitory activity against IL1RAP-associated cytokine signaling than CAN10 (Figure S4). Thus, DXP-006 (10 mg/kg) or LAD191 (10 mg/kg) was administered on days 0, 4, and 8 to evaluate the drug efficacy. (Figure 4A).

**Fig. 4.**
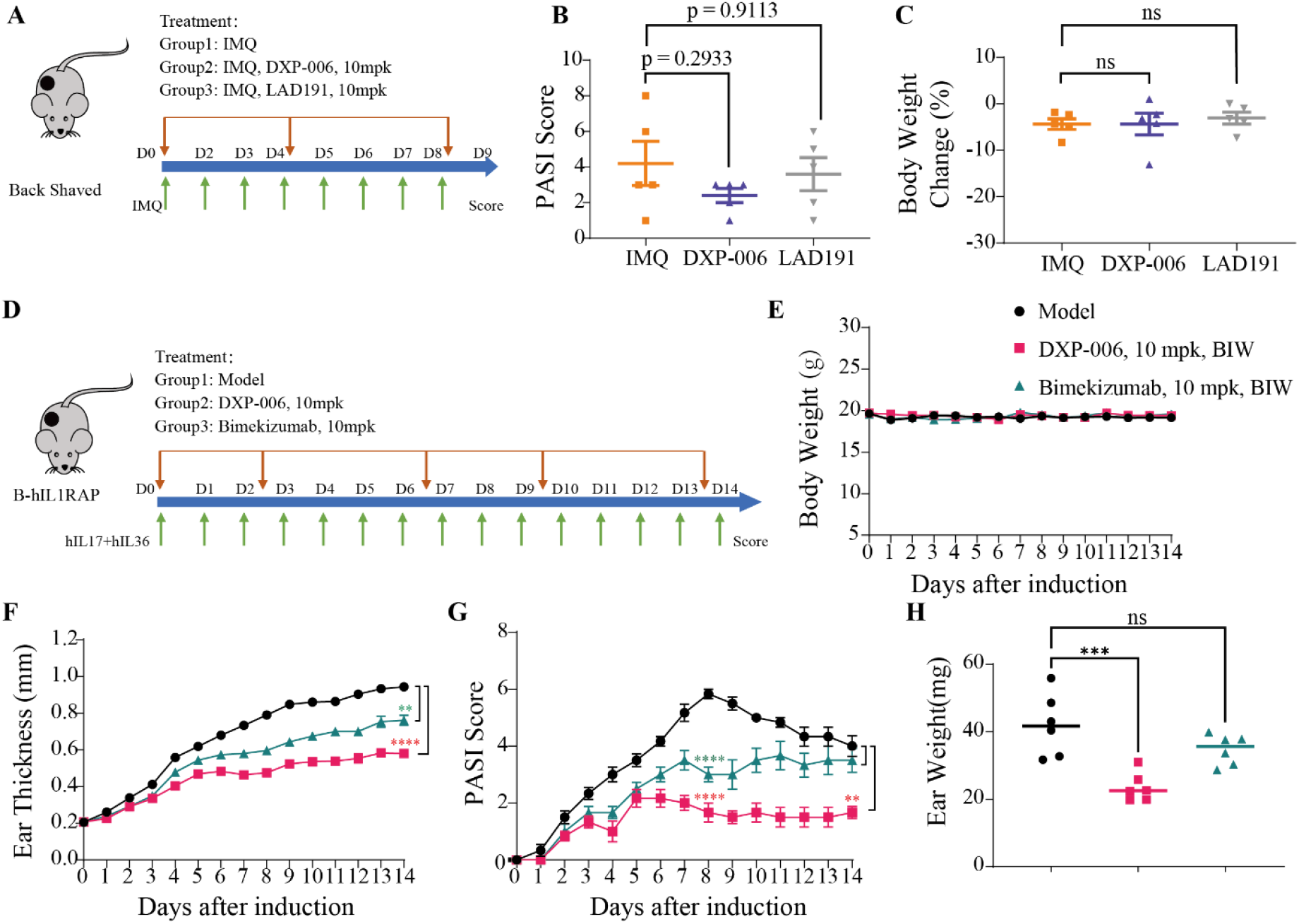
DXP-006 demonstrated superior efficacy in mouse psoriasis models. (A–C) In vivo efficacy of DXP-106 and Nadunolimab in a CDX mouse model bearing the human NSCLC cell line NCI-H358. Schematic illustration of the experimental design (A), with tumor volumes (B) and body weights (C) measured at the indicated time points. (D–H) In vivo efficacy of DXP-006 and in an IL-17- and IL-36-induced mouse model of psoriasis. Schematic illustration of the experimental design (D), with body weights (E), ear thickness (F), Clinical score (F) and ear weight (H) measured at the indicated time points. Statistical significance was defined as follows: *P < 0.05, **P < 0.01, ***P < 0.001, and ****P < 0.0001.

Compared with the untreated group, DXP-006 markedly alleviated skin lesions in the mouse back, as assessed by PASI scoring, and its efficacy surpassed that of LAD191 (Figure 4B). In terms of toxicity, neither DXP-006 nor LAD191 induced any significant weight loss in mice (Figure 4C). Therefore, DXP-006 effectively mitigates disease progression in IL-1 signaling–associated autoinflammatory disorders.

Further, the *in vivo* therapeutic effects of DXP-006 (10 mg/kg) or the IL-17 monoclonal antibody Bimekizumab (10 mg/kg) were assessed in IL-17- and IL-36-induced psoriasis models established in IL1RAP-humanized mice (Figure 4D). Throughout the experimental period, no significant changes in body weight were observed in any group, and all mice remained well-tolerated until the study endpoint (Figure 4E). Ear thickness measurements revealed that at Day 14 (study endpoint), compared to the model group, both DXP-006 and Bimekizumab treatment groups showed significant reductions in ear thicknes (Figure 4F). PASI scoring results demonstrated that at the disease peak (Day 8), compared to the model group, both DXP-006 and Bimekizumab treatment groups exhibited significantly reduced PASI scores. At the study endpoint (Day 14), compared to the model group, the DXP-006 group showed a significant reduction in PASI score, whereas the Bimekizumab group exhibited a decreasing trend without statistical significance (Figure 4G). Ear weight analysis indicated that at Day 14 (study endpoint), compared to the model group, the DXP-006 group showed a significant reduction in ear weight, while the Bimekizumab group displayed a decreasing trend without statistical significance (Figure 4H).

Collectively, these findings demonstrate that DXP-006, administered at 10 mg/kg via intraperitoneal injection twice weekly, conferred significant therapeutic benefits across multiple disease parameters in the psoriasis mouse model, including the suppression of ear thickening, the attenuation of clinical severity at peak disease, and the reduction of both clinical scores and ear eight at the study endpoint. Notably, the therapeutic efficacy of DXP-006 was significantly superior to that of Bimekizumab across all evaluated endpoints.

### Superior Efficacy of DXP-106 in Mouse Tumor Model

Since IL-1 signaling promotes tumor growth and metastasis, and to investigate the antitumor effects of blocking IL1RAP-mediated IL-1 signaling, we evaluated the *in vivo* efficacy of DXP-106 monotherapy in an NCI-H358 cell-derived xenograft (CDX) model. NCI-H358 cells highly express IL1RAP (Figure S5). CB17 SCID mice were inoculated with NCI-H358 cells (Figure 5A). From day 9 after inoculation, vehicles, Nadunolimab (10mg/kg, i.p, BIW) or DXP-106 (10mg/kg, i.p, BIW) were administered. The tumor volume and body weight were measured.

**Fig. 5.**
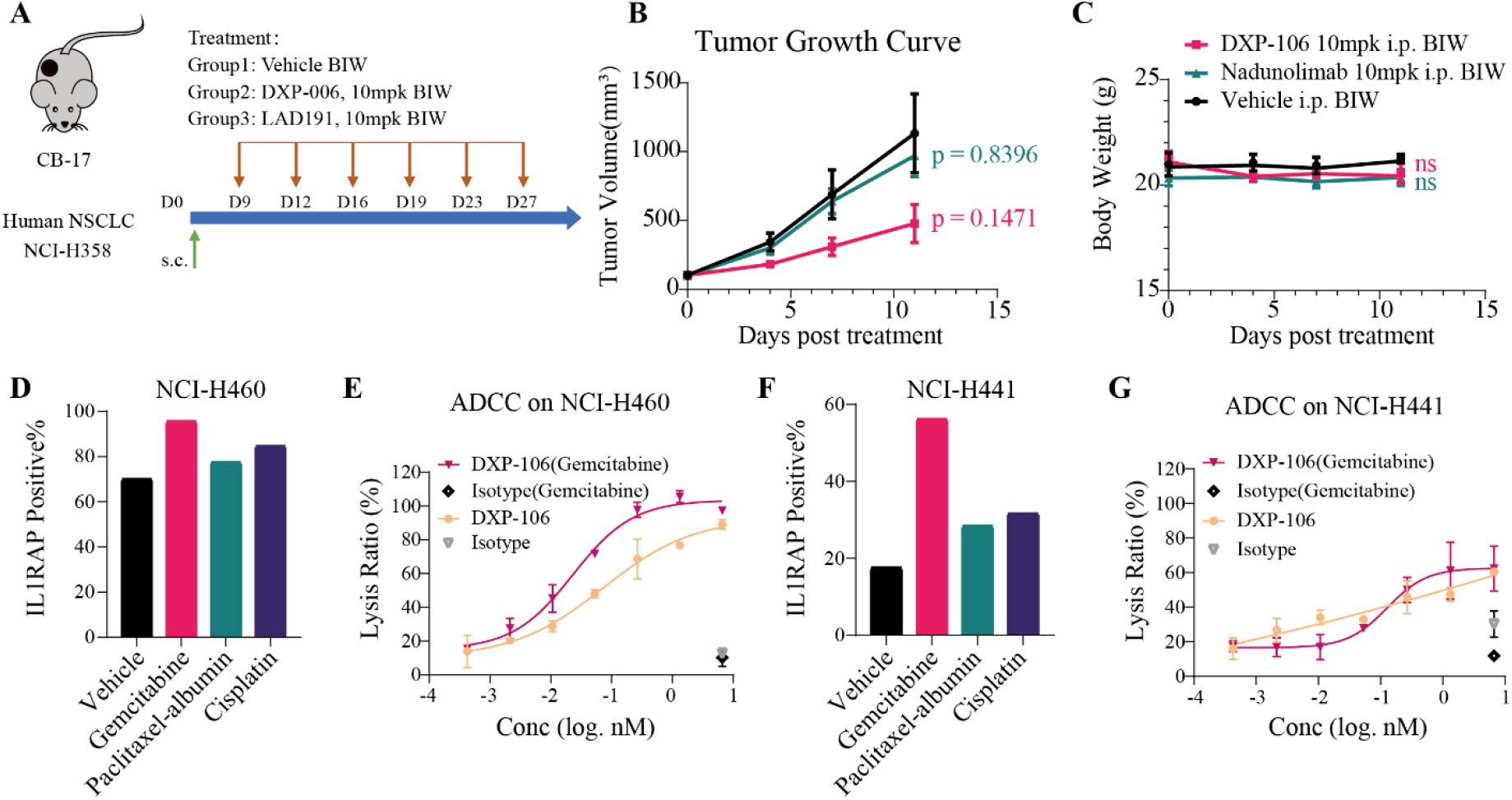
DXP-106 demonstrated superior in vitro and vivo anti-tumor efficacy. (A–C) In vivo efficacy of DXP-106 and Nadunolimab in a CDX mouse model bearing the human NSCLC cell line NCI-H358. Schematic illustration of the experimental design (A), with tumor volumes (B) and body weights (C) measured at the indicated time points. (D,F) FACS assessment of IL1RAP expression levels in NCI-H460 (D) and NCI-H441 (F) tumor cells after treatment with the chemotherapeutic agents, gemcitabine, paclitaxel-albumin, and cisplatin. (E,G) Assessment of DXP-106-mediated ADCC killing of NCI-H460 (E) and NCI-H441 (G) tumor cells before and after gemcitabine treatment. Statistical significance was defined as follows: *P < 0.05, **P < 0.01,***P < 0.001, and ****P < 0.0001.

Compared with both the vehicle-treated group and the Nadunolimab-treated group, tumor growth was significantly inhibited in mice that received DXP-106 treatment (Figure 5B and S6). No significant differences in body weight were observed among the DXP-106-treated, Nadunolimab-treated, and vehicle-treated groups (Figure 5C). DXP-106 monotherapy in NCI-H358 CDX model demonstrated significant in vivo efficacy.

IL1RAP has been found to be upregulated in tumor cells and the tumor microenvironment across a range of malignancies, such as acute myeloid leukemia, pancreatic ductal adenocarcinoma, and non-small cell lung cancer, among others. Chemotherapy results in increased IL1α levels within the tumor microenvironment, which further promotes resistance to chemotherapy (24).

We examined the expression levels of IL1RAP in various tumor cell lines following treatment with chemotherapy drugs, including cisplatin, paclitaxel-albumin, and gemcitabine (Figure 5D; 5F and S5). The results indicated that NCI-H460 (large-cell lung cancer) and NCI-H358 (non-small cell lung cancer) exhibited high expression of IL1RAP, irrespective of the treatment. Conversely, the expression of IL1RAP in HCT116 (colorectal cancer), CaPan-2 (pancreatic cancer), and NCI-H441 (papillary adenocarcinoma of the lung) was notably elevated after stimulation with gemcitabine. Additionally, the expression of IL1RAP in HCT116, CaPan-2, and EBC-1 (lung squamous cell carcinoma) increased following stimulation with cisplatin.

To determine if DXP-106 can induce antibody-dependent cell-mediated cytotoxicity (ADCC) against IL1RAP-expressing tumor cells, NCI-H460 cells with high IL1RAP expression and NCI-H441 cells with low IL1RAP expression were selected as target cells without gemcitabine treatment. Peripheral blood mononuclear cells (PBMCs) were used as effector cells to evaluate the antibody’s ADCC activity. As indicated, the NCI-H460 group demonstrated a higher overall ADCC activity of the antibody, and treatment with gemcitabine further increased this activity (Figure 5E). In contrast, the NCI-H441 group showed lower ADCC activity, though gemcitabine treatment slightly enhanced it (Figure 5G). Chemotherapy treatment augmented the ADCC effect of DXP-106, suggesting that combining chemotherapy with DXP-106 may overcome chemoresistance.

## DISCUSSION

In this study, we present the preclinical characterization of two novel anti-IL1RAP antibodies, DXP-006 and DXP-106, engineered to achieve broad and potent inhibition of the IL-1 cytokine family. Our structural and functional data reveal that by strategically targeting a critical functional epitope on the IL1RAP D2 domain, these antibodies effectively block signaling mediated by IL1α, IL1β, IL33, and all three IL-36 agonists, outperforming existing benchmark antibodies such as CAN10 and Nadunolimab in both affinity and functional potency.

The structural basis for this pan-inhibitory activity lies in the engagement of the c2d2 loop, a region essential for IL1RAP’s recruitment to multiple cytokine-receptor complexes. Cryo-EM analysis provided the first high-resolution view of this interaction, revealing that DXP-006 binds the c2d2 loop with high affinity via a previously uncharacterized binding mode, sterically hindering the formation of functional signaling assemblies. While the previously reported antibody CAN10 also targets the c2d2 loop, DXP-006 and DXP-106 exhibit superior binding kinetics and more efficient epitope “locking,” translating into consistently lower IC50 values across all cytokine pathways tested. Importantly, in models of combined cytokine stimulation—mimicking the complex milieu of human inflammatory diseases—single-target inhibitors showed limited efficacy, whereas DXP-006 and DXP-106 achieved near-complete suppression. This underscores the therapeutic advantage of co-receptor blockade in diseases such as psoriasis, rheumatoid arthritis, and certain solid tumors, where cytokine redundancy and synergy drive pathology.

Our *in vivo* studies further highlight the distinct therapeutic profiles of the two antibodies. DXP-006, optimized for extended half-life via Fc mutations (L234A/L235A/M252Y/S254T/T256E), demonstrated remarkable efficacy in psoriasis models, outperforming both an IL-17–targeted biologic and another IL1RAP antagonist. These results challenge the prevailing focus on downstream cytokines and suggest that upstream inhibition of IL-1 family signaling may provide a more comprehensive reset of the inflammatory microenvironment, particularly in conditions like generalized pustular psoriasis where IL-36 plays a dominant role. In diseases driven by epithelial-immune crosstalk (Psoriasis, IBD, Atopic Dermatitis), targeting the upstream “alarmin sensors” (IL-1, IL-33, IL-36) via IL1RAP may provide a more comprehensive “reset” of the tissue microenvironment. DXP-006 is particularly well-positioned for generalized pustular psoriasis (GPP) and palmoplantar pustulosis, conditions where IL-36 is the dominant driver.

DXP-106, designed with ADCC-enhancing Fc modifications, offers a multifaceted mechanism of action in oncology. It disrupts IL-1–driven tumor promotion and chemoresistance while actively engaging immune effector cells to eliminate IL1RAP-expressing tumor cells. The observed synergy with chemotherapy is particularly compelling, as chemotherapy-induced IL1RAP upversion creates a therapeutic window for enhanced ADCC-mediated clearance of residual, often resistant, tumor cells. This dual mechanism positions DXP-106 as a promising candidate for combination regimens, not only with chemotherapy but also with immune checkpoint inhibitors (PD-1/PD-L1 blockade). Since IL1β and IL33 are known to drive resistance to checkpoint blockade by recruiting MDSCs and Tregs (27), DXP-106 could serve as an ideal partner to “cleanse” the TME of immunosuppressive myeloid cells, thereby reinvigorating T-cell responses.

However, our study has several limitations that warrant discussion. While mouse models provide valuable insights, they do not fully recapitulate the complexity of human immune networks or the heterogeneity of tumor microenvironments. Additionally, the long-term safety of simultaneously blocking multiple cytokine pathways remains to be thoroughly evaluated. Future studies should include comprehensive toxicology and pharmacokinetic assessments in non-human primates to de-risk clinical translation. To translate these compelling preclinical findings into clinical applications, a defined path forward is required. For DXP-006, Phase Ib/IIa trials in patients with IL-1/IL-36-driven diseases such as psoriasis, hidradenitis suppurativa, or inflammatory bowel disease would be a logical starting point to demonstrate proof-of-concept for its broad anti-inflammatory activity. For DXP-106, initial clinical evaluation should focus on oncology, particularly in solid tumors known to overexpress IL1RAP and where chemotherapy is a standard of care (e.g., pancreatic ductal adenocarcinoma or non-small cell lung cancer). Furthermore, exploring combinations with immune checkpoint inhibitors represents a highly promising future direction, given the established role of IL-1 in modulating immunosuppression.

In conclusion, DXP-006 and DXP-106 represent an advancement in the targeting of IL-1 family signaling. By combining structural insights with rational antibody engineering, we have developed two high-affinity molecules capable of overcoming the limitations of single-cytokine inhibition. Their distinct designs cater to complementary therapeutic needs—broad anti-inflammatory activity for DXP-006 and immune-engaging antitumor efficacy for DXP-106—supporting their continued development as promising therapeutics for a range of inflammatory and malignant diseases.

## MATERIALS AND METHODS

### Study design

This study employed a multi-stage integrated research design encompassing antibody engineering, structural biology, biochemical characterization, and functional validation in both cellular and animal disease models.Rabbit-derived monoclonal antibodies targeting human IL1RAP domain 2 (D2) were isolated and subsequently humanized to generate DXP-006 and DXP-106, with distinct Fc engineering strategies—DXP-006 incorporating L234A/L235A/M252Y/S254T/T256E mutations for extended half-life and reduced effector function, and DXP-106 featuring a defucosylated Fc to enhance antibody-dependent cellular cytotoxicity (ADCC). The binding kinetics and epitope specificity were determined by Bio-Layer Interferometry and domain-specific ELISA, while the molecular mechanism of IL1RAP recognition was elucidated through cryo-electron microscopy (cryo-EM) structural analysis of the DXP-006/IL1RAP complex. Functional potency was assessed using NF-κB/AP-1 reporter assays in HEK293 cells for IL1α/IL1β pathway inhibition, IL-8 secretion assays in human umbilical vein endothelial cells (HUVEC) for IL33 blockade, and A431 cell-based assays for IL-36α/β/γ inhibition, with IC50 values benchmarked against clinical antibodies including CAN10, Nidanilimab, Canakinumab, Anakinra, and Spesolimab. Synergistic cytokine neutralization was evaluated using multi-cytokine stimulation models. For *in vivo* validation, therapeutic efficacy was tested in the imiquimod-induced psoriasis model in C57BL/6 mice and the IL-17/IL-36-driven ear inflammation model in IL1RAP-humanized mice, with disease severity assessed by PASI scoring, ear thickness measurements, and histopathological analysis. Antitumor activity was evaluated in the NCI-H358 cell-derived xenograft (CDX) model in CB17 SCID mice with tumor volume monitoring and body weight assessment. ADCC mechanisms were further investigated using PBMC-mediated cytotoxicity assays against chemotherapy-pretreated tumor cells expressing varying levels of IL1RAP. All animal experiments were randomized and blinded for outcome assessment, with sample sizes predetermined based on power analysis to ensure statistical rigor.

### Mice

CB-17 female mice were obtained from Beijing Vital River Laboratory Animal Technology Co., Ltd. B-hIL1RAP mice were obtained from Biocytogen Pharmaceuticals (Beijing) Co., Ltd. All mice were bred in specific pathogen-free conditions, housed in cages with five mice per cage, and kept on in a regular 12 h light/12 h dark cycle (lights on at 7:00 am). The temperature was 24 ± 2 °C and humidity was 40–70%.

### Cell lines

For cell signaling assay, HEK293/AP1-NFkB-LUC reporter cell line and epidermoid carcinoma cell line A431 were purchased from Sino Biological Inc. HEK293/AP1-NFkB-LUC cultured in MEM supplemented with 10% fetal bovine serum and 1% penicillin/streptomycin. A431 cultured in DMEM supplemented with 10% fetal bovine serum and 1% penicillin/streptomycin. Human Umbilical Vein Endothelial Cells (HUVEC) was purchased from Lifeline® Cell Technology (catalog no. FC-0003) and cultured in M199 supplemented with 10% fetal bovine serum and 5% L-Gln.

The human non-small cell lung cancer cell line NCI-H358, used to establish the mouse CDX model, was purchased from Wuhan Procell (catalog no. CL-0400) and cultured in RPMI-1640 supplemented with 10% fetal bovine serum and 1% penicillin/streptomycin.

For antibody ADCC activity detection, the human large-cell lung cancer cell line NCI-H460 was purchased from Wuhan Procell (catalog no. CL-0299) and the human lung adenocarcinoma cell line NCI-H441 was purchased from Nanjing Cobioer (catalog no. CBP60137). Both cultured in RPMI-1640 supplemented with 10% fetal bovine serum and 1% penicillin/streptomycin.

### Cell signaling assays

Cell signaling assays of IL1α and IL1β for DXP-006 and DXP-106 were performed using HEK293-AP1-NFkB-Luc −9 cells. For measuring the inhibition of IL1α or IL1β signaling, the appropriate HEK293 cells were seeded into 96-well plates and allowed to settle overnight before continuing. Cells were treated with escalating concentrations of DXP-006, DXP-106, or antagonists as indicated, followed by stimulation with IL-1α (Sinobiological, 10128-HNCH) or IL-1β (Sinobiological, 10139-HNAE) at their respective pre-determined EC_80_. After 6 h incubation at 37 °C, 5 % CO_2_, 25 µL of cell-lysis reagent was added to each well. The lysates were mixed thoroughly, and 20 µL from each well was transferred to a white 96-well plate. Following addition of 60 µL Luciferase Assay, luminescence was measured on a luminometer.

Cell signaling assays of IL33 or IL1α+IL1β+IL33 for DXP-006 and DXP-106 were performed using HUVEC. For measuring the inhibition of signaling, the appropriate HUVEC were seeded into 96-well plates and allowed to settle 4h at 37 °C, 5 % CO_2_ before continuing. Cells were treated with escalating concentrations of DXP-006, DXP-106, or antagonists as indicated, followed by stimulation with IL33 (Sinobiological, 10368-HNAE) or IL1α+IL1β+IL33 at their respective pre-determined EC_80_. After 24 h incubation at 37 °C, 5 % CO_2_, the culture supernatant was collected, and IL-8 levels were quantified by ELISA (Sinobiological, KIT10098).

Cell signaling assays of IL36α, IL36β, IL36γ or IL1α+IL1β+ IL36α+IL36β+IL36γ for DXP-006 and DXP-106 were performed using A431 cells. For measuring the inhibition of signaling, the appropriate A431 cells were seeded into 96-well plates and allowed to settle 4h at 37 °C, 5 % CO_2_ before continuing. Cells were treated with escalating concentrations of DXP-006, DXP-106, or antagonists as indicated, followed by stimulation with IL36α (Sinobiological, 10607-HNCE1), IL36β (Sinobiological, 10579-HNCE1), IL36γ (Sinobiological, 10124-HNCE1) or IL1α+IL1β+ IL36α+IL36β+IL36γ at their respective pre-determined EC_80_. After 24 h incubation at 37 °C, 5 % CO_2_, the culture supernatant was collected, and IL-8 levels were quantified by ELISA (Sinobiological, KIT10098).

### Cryo-EM sample preparation and data collection and model building

For the preparation of the complexes, the purified IL1RAP was mixed with a 1.5-fold molar excess of fab DXP006 and another fab binding to D3 domain, incubated on ice for 40 min and injected onto a Superdex 200 Increase 5/150 column (Cytiva) equilibrated with buffer 1x PBS. SDS-PAGE analysis confirmed the formation of the DXP006-IL1RAP complexes in a stoichiometric ratio.

An aliquot of 4 μL protein sample of complex at a protein concentration of 0.5 mg/mL was loaded onto a glow-discharged 300 mesh grid (Quantifoil Au R1.2/1.3). The grids were blotted with a filter paper at 4 ° C and 100% humidity, and flash-cooled in liquid ethane using a Thermo Fisher Vitrobot Mark IV and screened using a 200 KV Talos Aectica.

Cryo-EM datasets were collected on a 300kV Thermo Fisher Titan Krios G4 electron microscope equipped with a Falcon 4 camera and a selectris X energy filter (GIF: a slit width of 10eV). The micrographs were collected at a calibrated magnification of x130,000 using the EPU software (Thermo Fisher Scientific), yielding a pixel size of 0.95 Å at object scale. Movies were recorded at an accumulated electron dose of 60e-Å-2 s-1 on each micrograph that was fractionated into a stack of 40 frames with a defocus range of −1.0 μm to ™2.0 μm.

Data processing was performed using cryoSPARC (v4.7.1). The data underwent several steps including Motion Correction, CTF Estimation, Create Templates, Template Picker, Extract from Micrographs, 2D classification, 2D selection for Ab-initio Reconstruction, and subsequent Homogeneous Refinement. Structural modeling and refinement were performed using WinCoot (v0.9.4.1) and Phenix (v1.21.2). Figures were generated using UCSF ChimeraX (v1.7).

### Enzyme-Linked Immunosorbent Assay (ELISA)

The binding specificity of the purified antibodies to different domains of IL1RAP was assessed by ELISA.

Microtiter plates were coated overnight at 4°C with 100 µL per well of recombinant human IL1RAP proteins at concentrations of 0.1 µg/mL and 1 µg/mL. The proteins included full-length IL1RAP-Fc, IL1RAP Domain 1 (D1), Domains 1 & 2 (D1&D2), and IL1RAP Domain 3 (D3). Following the coating step, the plates were emptied and tapped dry. Non-specific binding sites were blocked by adding 300 µL per well of 2% (w/v) BSA in PBS and incubating for 1 hour at room temperature with sealing. The plates were washed twice with 300 µL per well of wash buffer and thoroughly dried after the final wash.

Purified antibodies were diluted to a working concentration of 1 µg/mL in sample dilution buffer. For the titration curve, antibodies were serially diluted using a three-fold dilution series starting from 60 nM, generating seven concentration points. The plates were washed three times. Diluted antibodies (100 µL per well) were added to the plates, which were then incubated for 2 hours at room temperature. Then, 100 µL per well of horseradish peroxidase (HRP)-conjugated Goat Anti-Human IgG F(ab’)_2_ fragment was added, and the plates were incubated for 1 hour at room temperature. 200 µL of the mixed tetramethylbenzidine (TMB) solution was added to each well, followed by incubation for 20 minutes at room temperature in the dark. The enzymatic reaction was stopped by adding 50 µL of stop solution per well. The optical density (OD) at 450 nm was measured immediately using a microplate reader.

### Binding Affinity Measurement by Bio-Layer Interferometry (BLI)

The antigen was serially diluted in QBuffer (PBS supplemented with 0.02% Tween-20 and 0.2% IgG-free BSA). A starting concentration of 200 nM was subjected to two-fold serial dilutions to generate a total of seven concentration gradients for the analysis. All antibody samples were diluted to a uniform concentration of 10 µg/mL using QBuffer. A volume of 200 µL per sample was required for each analysis cycle, with a total of 2 mL prepared per antibody to accommodate the multi-concentration antigen measurement.

Following the software setup, the prepared samples and reagents were loaded into the designated wells of the sample plate and the Gator MAX plate as per the protocol. Protein A biosensors (Gator Bio, USA) were placed into the corresponding columns of the MAX plate for antibody capture. The assay procedure, including steps for sensor hydration, antibody loading, baseline establishment, association (antigen binding), and dissociation, was defined within the Gator Prime software. The run groups and timing parameters were configured accordingly before initiating the automated sequence.

The binding kinetics data was processed and analyzed using the integrated Gator Prime software. The sensorgrams were fitted to a 1:1 binding model to generate the binding curves. The equilibrium dissociation constant (KD), along with the coefficient of determination (Full R^2^) for the curve fit, were automatically calculated by the software to evaluate the binding affinity.

### Antibody-Dependent Cell-mediated Cytotoxicity (ADCC) Assay

Human lung carcinoma cell lines H460 and H441 were used as target cells. The cells were harvested, resuspended in RPMI-1640 medium, and counted. The cell density was adjusted to 3 × 10^5 cells/mL. Subsequently, 1 mL of the cell suspension was seeded into each well of a 6-well plate. The plate was incubated overnight at 37°C in a 5% CO2 humidified incubator to allow cell adherence. Following incubation, the cells were treated with 1 mL per well of 10 nM Gemcitabine (prepared in RPMI-1640) or vehicle control and incubated for an additional 24 hours under the same conditions.

Peripheral Blood Mononuclear Cells (PBMCs) were thawed, resuspended in RPMI-1640 medium, counted, and incubated overnight at 37°C with 5% CO2 to allow recovery before being used as effector cells.

Gemcitabine-treated or untreated H460 and H441 target cells were collected, resuspended, and counted. The cell density was adjusted to 1 × 10^6 cells/mL. The cells were then labeled with the DELFIA® BATDA reagent at a ratio of 2.0 µL per million cells per mL of cell suspension. Labeling was performed by incubating the cells for 25 minutes at 37°C with 5% CO2. After incubation, the cells were washed three times with 3 mL of PBS (supplemented with 20 mM HEPES and 2 mM probenecid) by centrifugation at 1200 rpm for 4 minutes at room temperature. The labeled cells were finally resuspended in RPMI-1640 medium containing 2 mM probenecid at a density of 2 × 10^5 cells/mL. For the assay, 50 µL of this cell suspension was dispensed into each well of the assay plate. The target cells were then pre-incubated with DXP-106. Specifically, the antibody was diluted to the indicated concentrations, and 50 µL was added to the corresponding wells containing the labeled target cells. The plate was incubated for 15 minutes at 37°C with 5% CO2.

PBMC effector cells were collected and adjusted to a density of 3 × 10^6 cells/mL. A volume of 100 µL of this effector cell suspension was added to the wells containing the pre-incubated target cells, resulting in an Effector to Target (E:T) ratio of 30:1. The co-culture plate was then incubated for 4 hours at 37°C in a 5% CO2 atmosphere to allow ADCC to occur. Following the 4-hour co-incubation, the assay plate was centrifuged. From each well, 20 µL of the supernatant was carefully transferred to a dedicated detection plate. To this supernatant, 200 µL of the DELFIA® Europium solution was added per well. The detection plate was placed on an orbital shaker and agitated at 250 rpm for 15 minutes at room temperature to allow chelate formation. The time-resolved fluorescence was measured using a microplate fluorometer. The measurement parameters were an excitation wavelength of 340 nm and an emission wavelength of 615 nm, with a time delay of 400 µs and a measurement window of 400 µs.

### Evaluation of the Anti-tumor Efficacy of DXP-106 in an NCI-H358 CDX Model

The human lung cancer cell line NCI-H358 was used to establish the subcutaneous xenograft model. Female CB17 SCID mice, aged 6-8 weeks, were housed under specific pathogen-free conditions. NCI-H358 cells were harvested during the logarithmic growth phase and resuspended in PBS. Approximately 5 x 10^6 cells in a 100 μL volume were inoculated subcutaneously into the right flank of each mouse.

When the average tumor volume reached approximately 50-100 mm^3^, the tumor-bearing mice were randomized into 3 groups (n=5 per group) based on tumor volume and body weight to ensure equivalent starting conditions across groups. Treatment was initiated immediately after randomization. The administered antibodys were dissolved in PBS. All treatments were administered via subcutaneous (s.c.) injection, twice per week. Body weight and tumor volume were measured twice weekly. Tumor volume was calculated using the standard formula: Volume (mm^3^) = (Length × Width^2^) / 2, where length is the longest dimension and width is the perpendicular shorter dimension. Mice were humanely euthanized when tumor volume exceeded 1500 mm^3^ or body weight loss surpassed 20% of the initial weight.

Datas were analyzed using one-way ANOVA for single-factor comparisons or two-way ANOVA for two-factor comparisons. Statistical significance was defined as follows: *P < 0.05, **P < 0.01,***P < 0.001, and ****P < 0.0001.

### Induction of Psoriasis-like Dermatitis Model and Treatment Protocol

A total of fifteen female C57BL/6 mice were used in the Imiquimod (IMQ)-induced mouse model of psoriasis. The mice were randomly assigned into three groups (n=5 per group) based on body weight, with the day of grouping designated as Day 0. Two days prior to the start of the experiment (Day -2), the dorsal skin of all mice was shaved to create an area of approximately 2 cm × 3 cm. The skin condition in the shaved area was observed to ensure it was intact before model induction. To induce a psoriasis-like skin inflammation, a daily topical dose of 62.5 mg of Imiquimod (IMQ) cream was applied uniformly to the shaved back of each mouse from Day 0 to Day 7 (for 8 consecutive days).

A total of fifteen female B-hIL1RAP mice included in this study for the establishment of an IL-17- and IL-36-induced mouse model of psoriasis. The mice were randomly assigned into three groups (n=5 per group) based on body weight, with the day of grouping designated as Day 0. Psoriasis-like skin inflammation was induced in mice by daily intradermal injection of 1 μg IL-17 and 1 μg IL-36 into the ear pinnae from Day 0 to Day 14 (total of 15 injections). Beginning on Day 0, test antibodys (DXP-006 or Bimekizumab) were administered twice weekly via intraperitoneal injection. Therapeutic efficacy was evaluated through multiple parameters: (1) daily measurement of ear thickness; (2) daily clinical observation and photographic documentation of ear lesions; (3) clinical scoring based on Psoriasis Area and Severity Index (PASI) criteria; and (4) ear weight at the study endpoint.

Datas were analyzed using one-way ANOVA for single-factor comparisons or two-way ANOVA for two-factor comparisons. Statistical significance was defined as follows: *P < 0.05, **P < 0.01, ***P < 0.001, and ****P < 0.0001.

### Statistical analysis

Data from mouse model of psoriasis and NCI-H358 CDX model were analyzed using one-way ANOVA for single-factor comparisons or two-way ANOVA for two-factor comparisons. Statistical significance was defined as follows: *P < 0.05, **P < 0.01, ***P < 0.001, and ****P < 0.0001.

## List of Supplementary Materials

Fig S1 to S6

Tables S1 to S2

## Author contributions

Conceptualization: Qinghao Liu, Qian Shi, Yunlong Cao

Methodology: Qinghao Liu, Yanxia Wang, Qian Shi, Yunlong Cao

Investigation: Yanxia Wang, Yaqi Ru, Qian Zhao, Zhen Zhang, Zhiqiang Li

Visualization: Qinghao Liu, Yanxia Wang

Funding acquisition: Qinghao Liu, Qian Shi, Yunlong Cao

Project administration: Qinghao Liu

Supervision: Qinghao Liu, Qian Shi, Yunlong Cao

Writing – original draft: Yan Jiang

Writing – review & editing: Yan Jiang, Qinghao Liu, Yunlong Cao

## Data and materials availability

Cryo-EM data for the DXP-006/IL1RAP complex have been deposited in the Protein Data Bank (PDB), with accession code 23QE, and in the Electron Microscopy Data Bank (EMDB), with accession code EMD-69166. Additional materials and data are available from the lead corresponding author (Y.C.) upon request and are subject to a Material and Data Transfer Agreement. All enquiries will be replied to within 7 working days. Source data are provided with this paper

‘n.t’. = not test

## Supplementary Materials

### Supplementary Figures

**Fig. S1.**
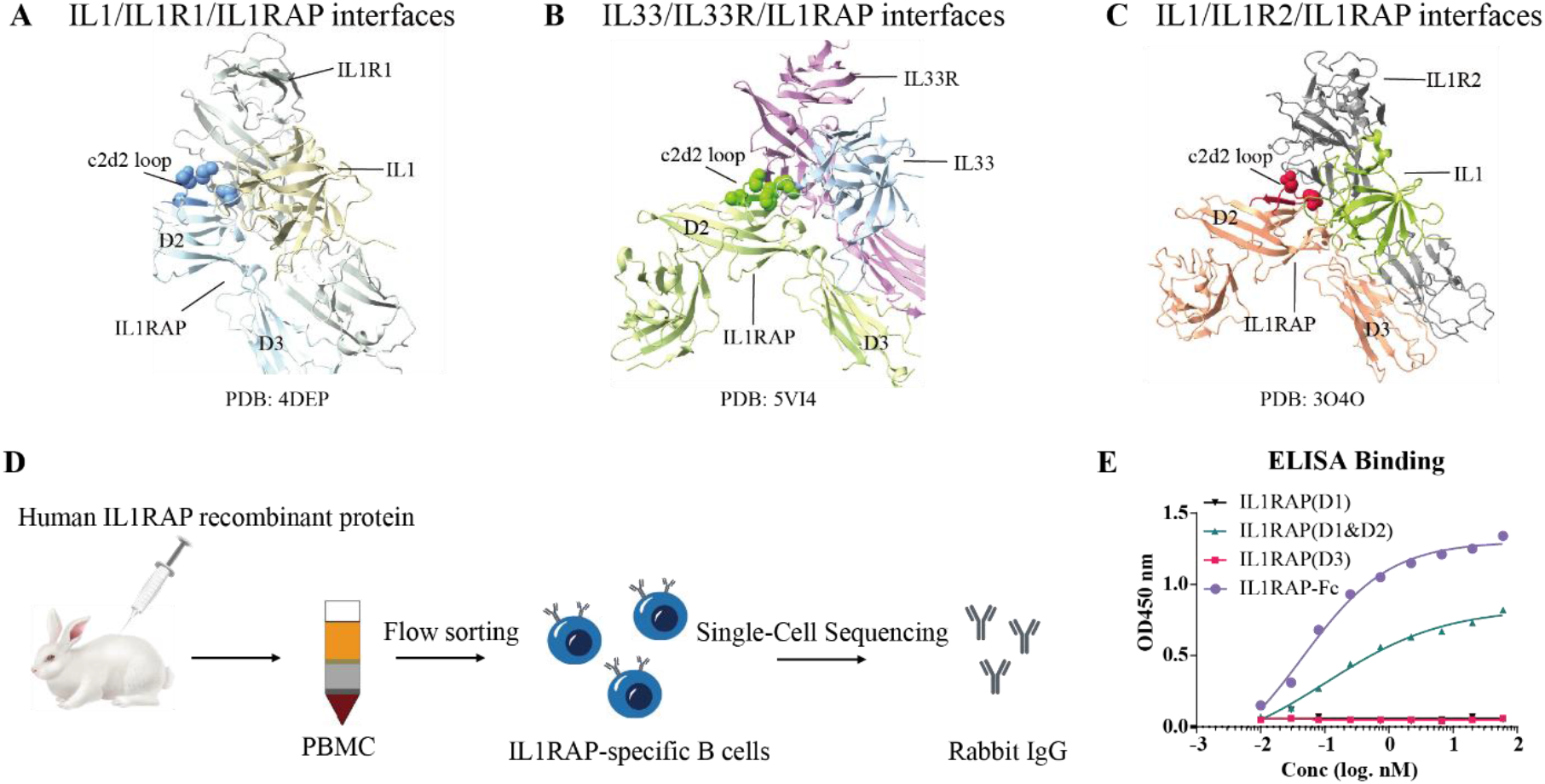
Antibodies that bind to domain 2 of IL1RAP were obtained by sequencing antigen-specific B cells from immunized rabbits. (A) Crystal structure of IL1RAP (PDB: 4DEP), with IL1 (olive), IL-1RI (grey) and c2d2 loop (blue) interfaces colored. (B) Crystal structure of IL1RAP (PDB: 5VI4), with IL33 (cyan), IL-33R (pink) and c2d2 loop (green) interfaces colored. (C) Crystal structure of IL1RAP (PDB: 3O4O), with IL1 (dark goldenrod), IL-1R2 (deep pink) and c2d2 loop (red) interfaces colored. (D) Schematic diagram of antibody discovery and screening workflow. (E) ELISA binding assessment of the parental antibody to full-length IL1RAP and its individual domains.

**Fig. S2.**
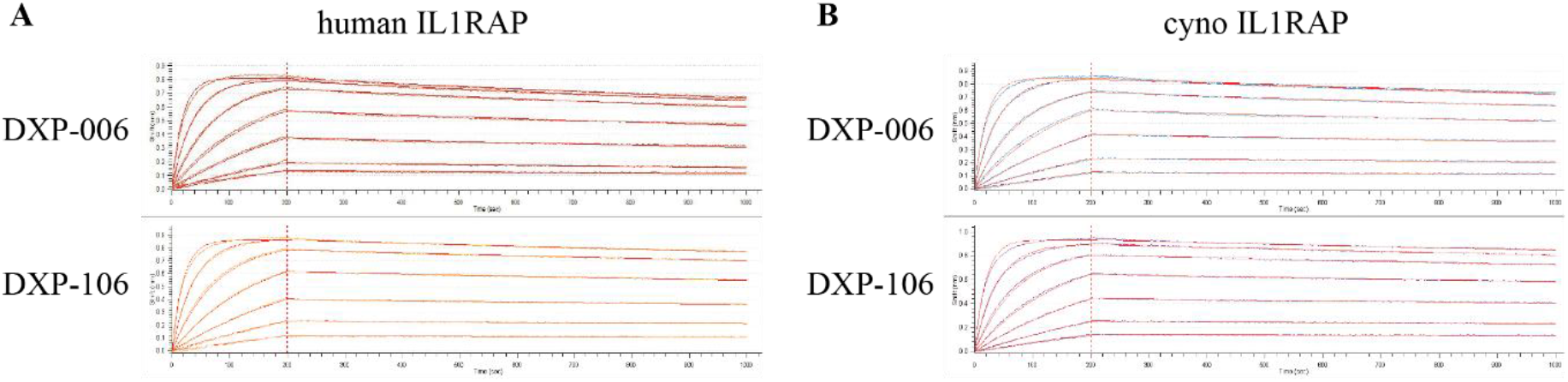
BLI analysis of DXP-006 and DXP-106 to human and cynomolgus IL1RAP. (A) BLI analysis of DXP-006 and DXP-106 to human IL1RAP. (B) BLI analysis of DXP-006 and DXP-106 to cynomolgus IL1RAP.

**Fig. S3.**
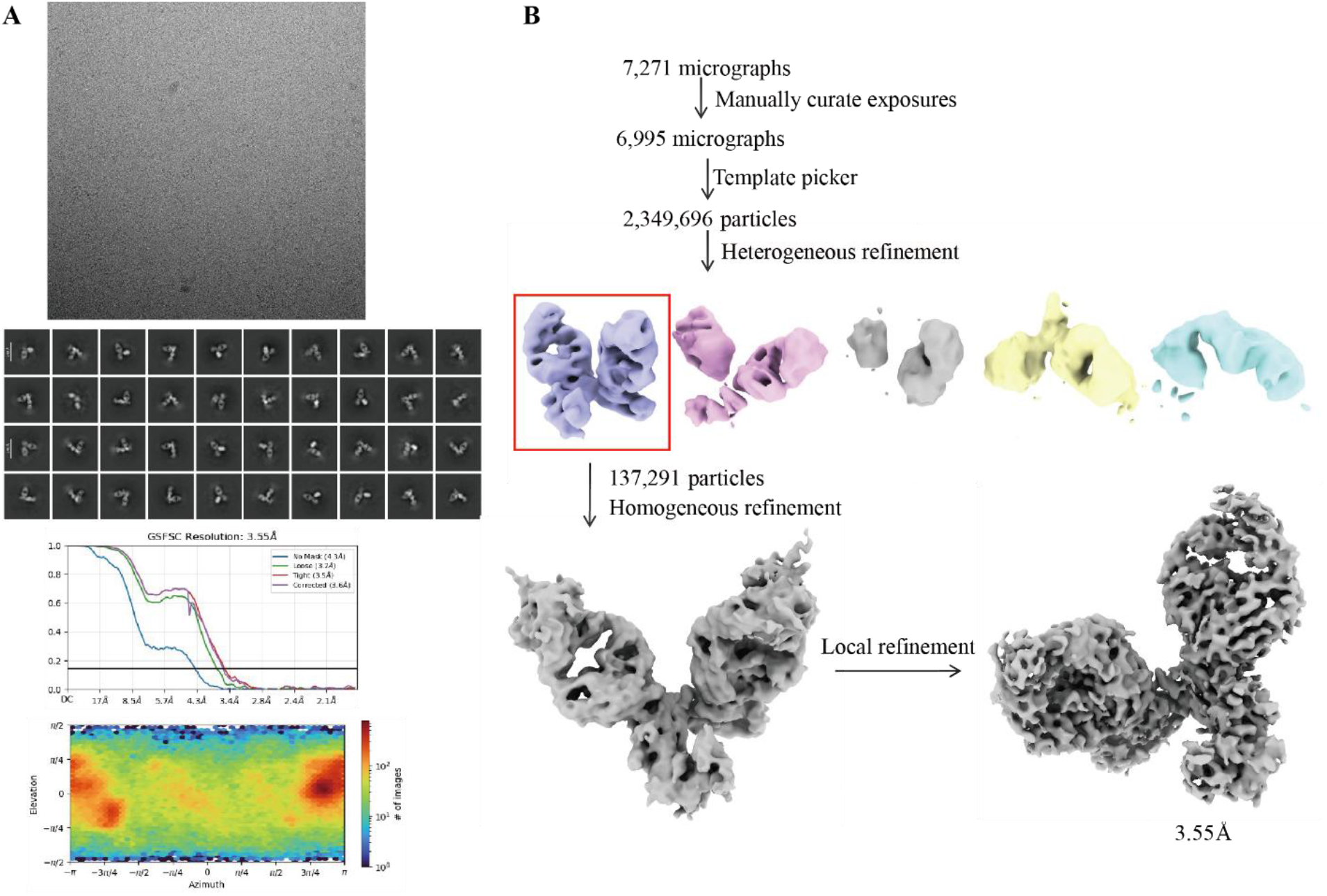
Workflow of the cryo-EM structure determination of the DXP-006–IL1RAP complex. (A) Representative cryo-EM micrograph of the DXP-006 Fab–IL1RAP complex acquired on a Titan Krios G4 microscope equipped with a Falcon 4 camera and Selectris X energy filter. (B) 2D class averages showing distinct secondary structural features and particle orientations.

**Fig. S4.**
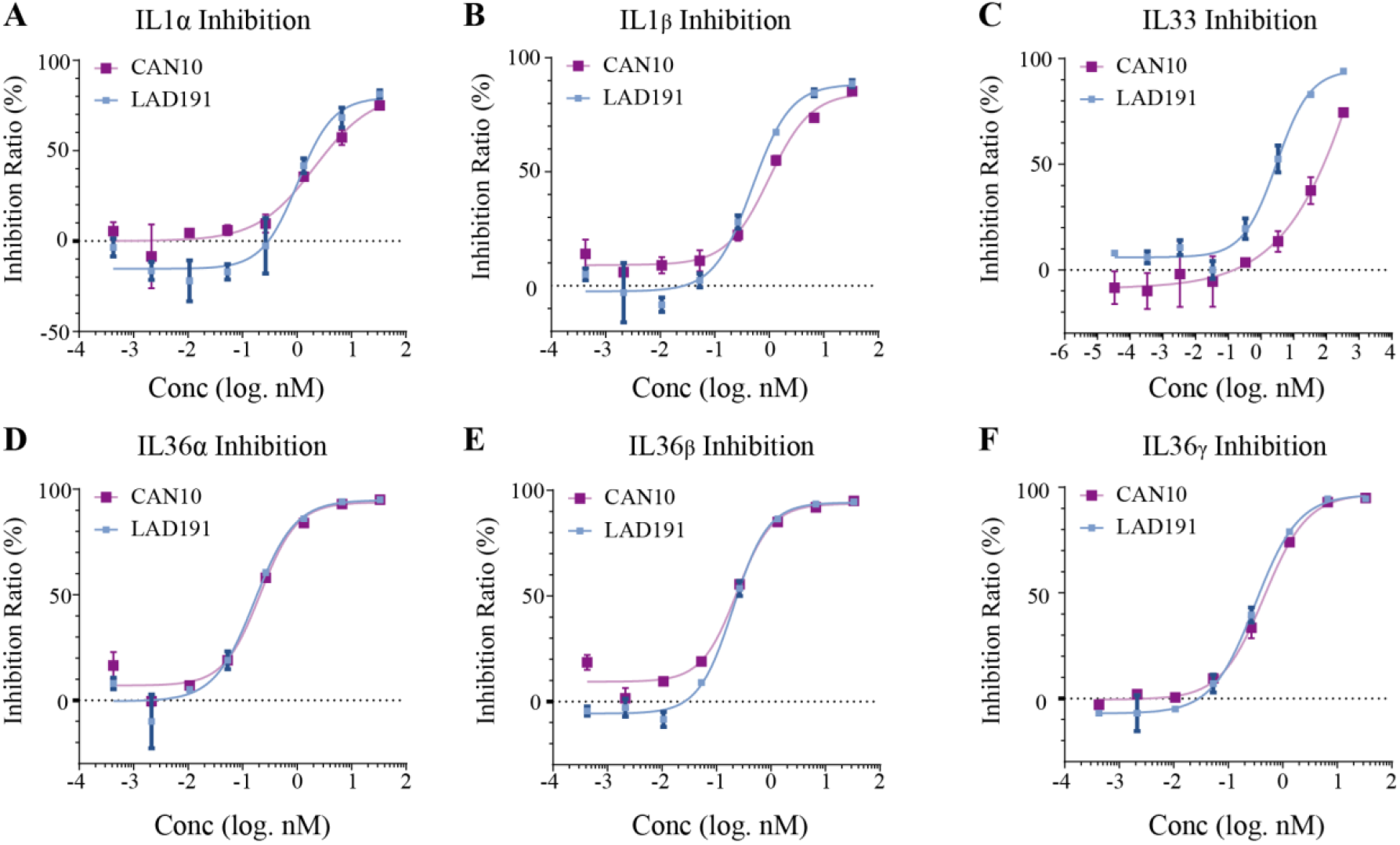
LAD191 demonstrated higher inhibitory potency compared to CAN10. (A-B) Inhibition of IL1α (A) and IL1β (B) induced signaling by indicated antibodies in HEK-Blue IL-1 reporter cells. (C) Inhibition of IL33 induced signaling by indicated antibodies in HUVEC cells. (D-F) Inhibition of IL36α (D), IL36β (E) and IL36γ (F) induced signaling by indicated antibodies in A431 cells.

**Fig. S5.**
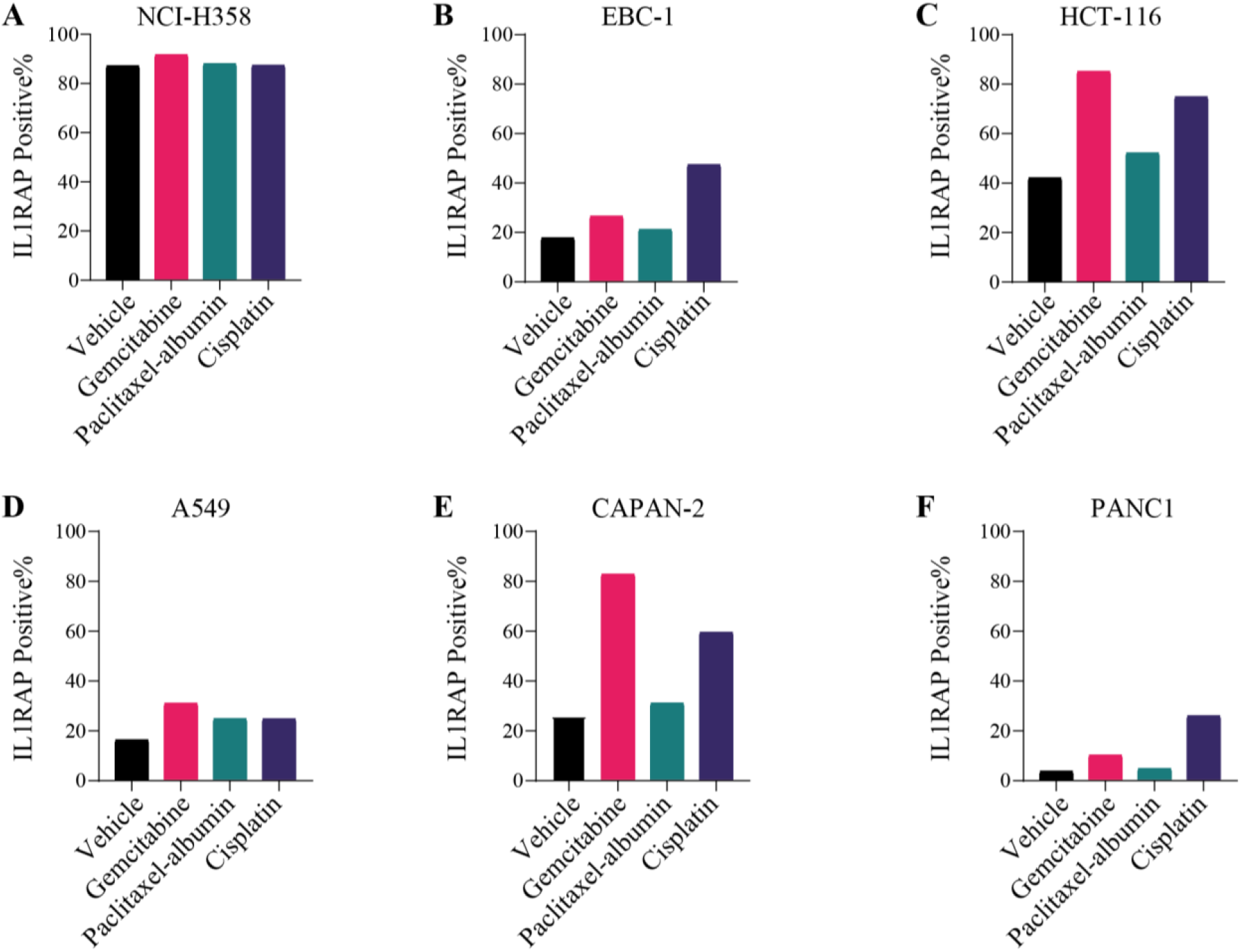
IL1RAP levels in tumor cells after treatment with the chemotherapeutic agents. (A-F) Flow cytometric analysis of IL1RAP surface expression on human cancer cell lines NCI-H358 (A), EBC-1 (B), HCT116 (C), A549 (D), CaPan-2 (3) and PANC1 (F) after 24-hour exposure to gemcitabine, paclitaxel-albumin, or cisplatin.

**Fig. S6.**
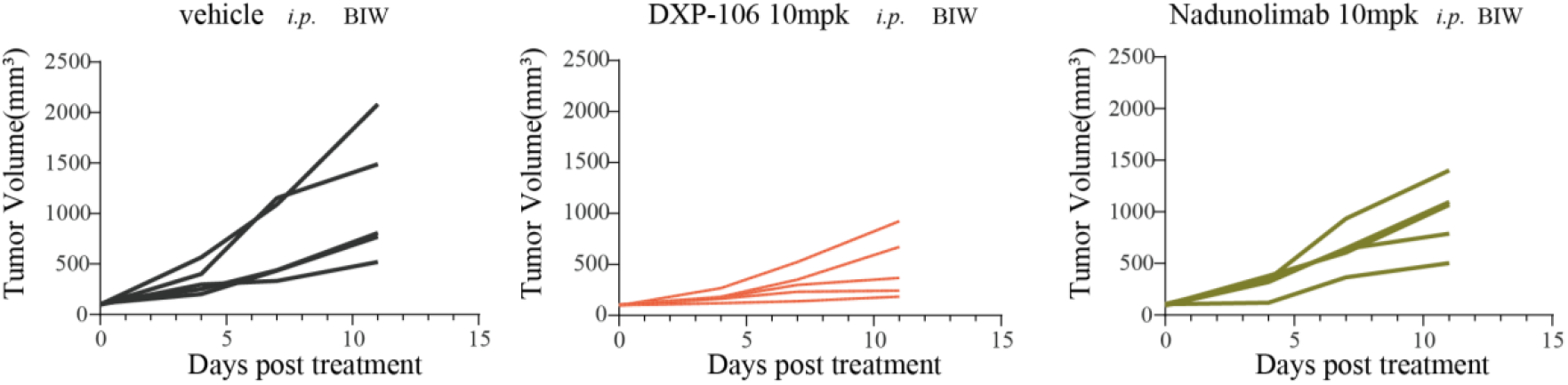
Individual tumor volumes in vehicle-treated, DXP-106-treated, and Nadunolimab-treated mice. Individual tumor growth curves of CB17 SCID mice bearing NCI-H358 subcutaneous xenografts. Mice were administered vehicle (PBS), Nadunolimab (10 mg/kg), or DXP-106 (10 mg/kg) twice weekly via intraperitoneal injection.

### Supplementary Tables

**Table S1.**
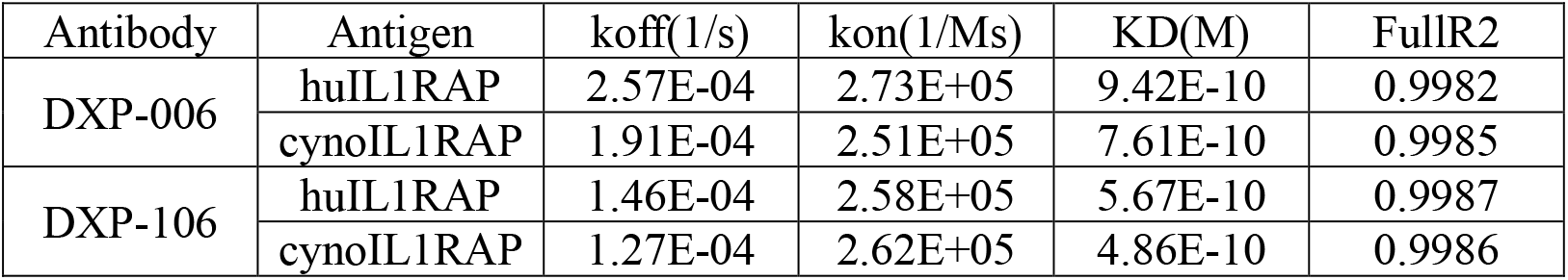
The binding affinity of DXP-006 and DXP-106 to human and cynomolgus IL1RAP.

**Table S2.**
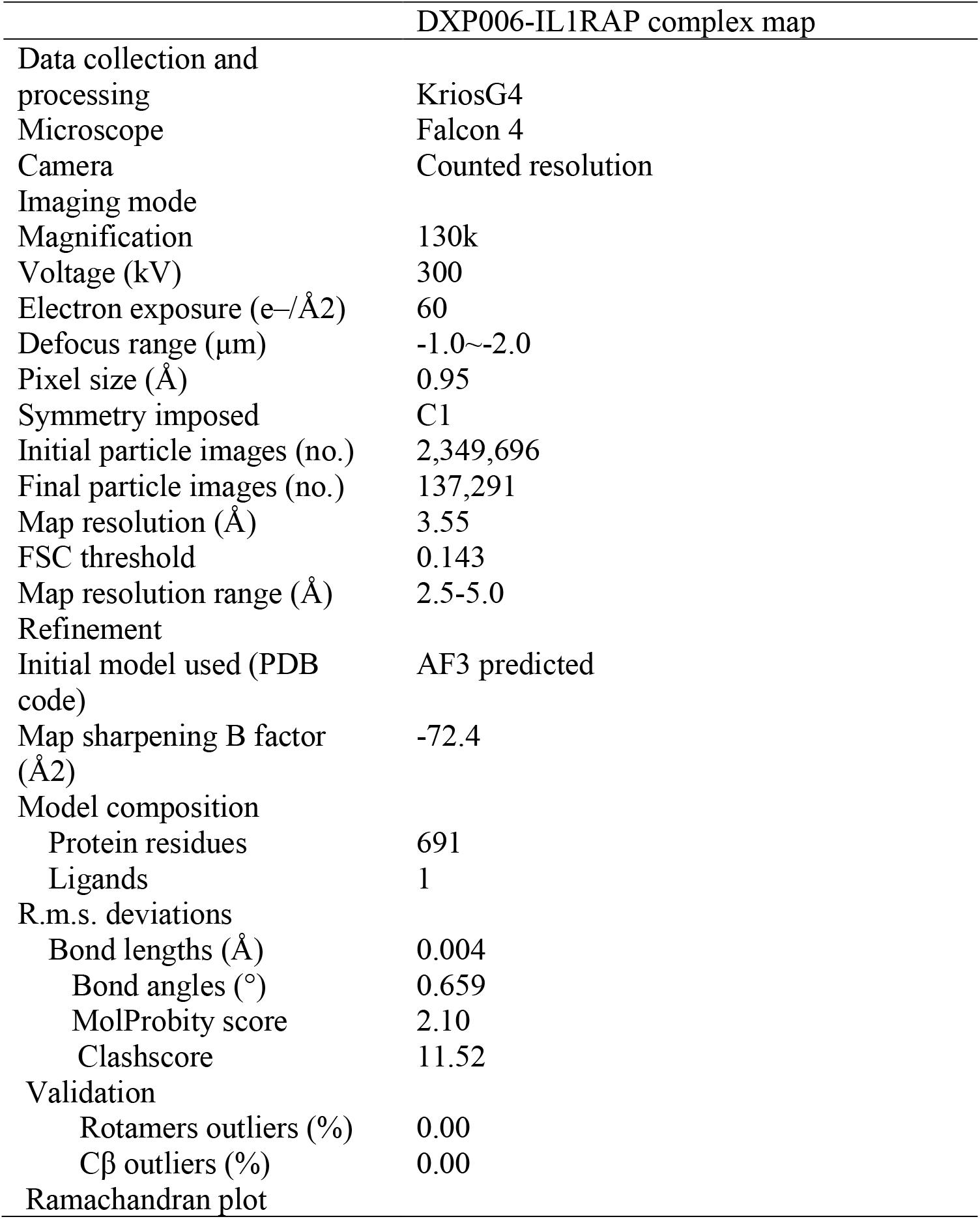
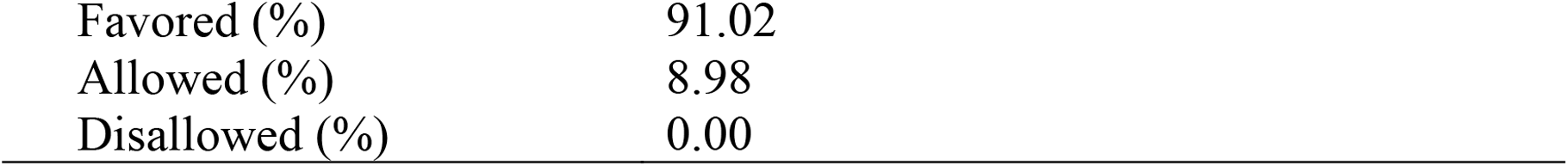
Cryo-EM data collection, refinement, and validation statistics.

| DXP006-IL1RAP complex map |  |
| --- | --- |
| Data collection and processing | KriosG4 |
| Microscope | Falcon 4 |
| Camera | Counted resolution |
| Imaging mode |  |
| Magnification | 130k |
| Voltage (kV) | 300 |
| Electron exposure (e-/Å<sup>2</sup>) | 60 |
| Defocus range (µm) | -1.0~-2.0 |
| Pixel size (Å) | 0.95 |
| Symmetry imposed | C1 |
| Initial particle images (no.) | 2,349,696 |
| Final particle images (no.) | 137,291 |
| Map resolution (Å) | 3.55 |
| FSC threshold | 0.143 |
| Map resolution range (Å) | 2.5-5.0 |
| Refinement |  |
| Initial model used (PDB code) | AF3 predicted |
| Map sharpening B factor (Å<sup>2</sup>) | -72.4 |
| Model composition |  |
| Protein residues | 691 |
| Ligands | 1 |
| R.m.s. deviations |  |
| Bond lengths (Å) | 0.004 |
| Bond angles (°) | 0.659 |
| MolProbity score | 2.10 |
| Clashscore | 11.52 |
| Validation |  |
| Rotamers outliers (%) | 0.00 |
| Cβ outliers (%) | 0.00 |
| Ramachandran plot |  |
| Favored (%) | 91.02 |
| Allowed (%) | 8.98 |
| Disallowed (%) | 0.00 |

## References and Notes

1. Sims, J., Smith, D. The IL-1 family: regulators of immunity. Nat. Rev. Immunol. 10, 89–102 (2010).

2. Garlanda C, Dinarello CA, Mantovani A. The interleukin-1 family: back to the future. Immunity 39(6), 1003–1018 (2013).

3. Chen Z, Bozec A, Ramming A, Schett G. Anti-inflammatory and immune-regulatory cytokines in rheumatoid arthritis. Nat. Rev. Rheumatol. 15, 9–17 (2019).

4. Liew FY, Girard JP, Turnquist HR. Interleukin-33 in health and disease. Nat. Rev. Immunol. 16(11), 676–689 (2016).

5. Boutet MA, Nerviani A, Pitzalis C. IL-36, IL-37, and IL-38 cytokines in skin and joint disease: A comprehensive review of their therapeutic potential. Int. J. Mol. Sci. 20(6), 1257 (2019).

6. Iznardo H, Puig L. The interleukin-1 family cytokines in psoriasis: pathogenetic role and therapeutic perspectives. Expert. Rev. Clin. Immunol. 17(2), 187–199 (2021).

7. Gabay C, Lamacchia C, Palmer G IL-1 pathways in inflammation and human diseases. Nat. Rev. Rheumatol 6(4), 232–241 (2010).

8. Zhou L, Todorovic V. Interleukin-36: Structure, Signaling and Function. Adv. Exp. Med. Biol.21, 191–210 (2021).

9. Fields JK, Günther S, Sundberg EJ. Structural Basis of IL-1 Family Cytokine Signaling. Front. Immunol. 20(10), 1412 (2019).

10. Thomas C, Bazan JF, Garcia KC. Structure of the activating IL-1 receptor signaling complex. Nat. Struct. Mol. Bio.l 19(4), 455–7 (2012).

11. Günther S, Deredge D, Bowers AL, Luchini A, Bonsor DA, Beadenkopf R, Liotta L, Wintrode PL, Sundberg EJ. IL-1 Family Cytokines Use Distinct Molecular Mechanisms to Signal through Their Shared Co-receptor. Immunity 19, 47(3), 510-523.e4 (2017).

12. Calabrese L, Fiocco Z, Satoh TK, Peris K, French LE. Therapeutic potential of targeting interleukin-1 family cytokines in chronic inflammatory skin diseases. Br. J. Dermatol. 186(6), 925–941 (2022).

13. Zarezadeh Mehrabadi A, Aghamohamadi N, Khoshmirsafa M, Aghamajidi A, Pilehforoshha M, Massoumi R, Falak R. The roles of interleukin-1 receptor accessory protein in certain inflammatory conditions. Immunology 166(1), 38–46 (2022).

14. Frenay J, Bellaye PS, Oudot A, Helbling A, Petitot C, Ferrand C, Collin B, Dias AMM. IL-1RAP, a Key Therapeutic Target in Cancer. International Journal of Molecular Sciences 23(23), 14918 (2022).

15. De Boer B, Sheveleva S, Apelt K, Vellenga E, Mulder AB, Huls G, Jacob Schuringa J. The IL1-IL1RAP axis plays an important role in the inflammatory leukemic niche that favors acute myeloid leukemia proliferation over normal hematopoiesis. Haematologica 106(12), 3067–3078 (2021).

16. Garlanda C, Mantovani A. Interleukin-1 in tumor progression, therapy, and prevention. Cancer Cell 39(8), 1023–1027 (2021).

17. Apte RN, Dotan S, Elkabets M, White MR, Reich E, Carmi Y, Song X, Dvozkin T, Krelin Y, Voronov E. The involvement of IL-1 in tumorigenesis, tumor invasiveness, metastasis and tumor-host interactions. Cancer Metastasis. Rev. 25, 387–408 (2006).

18. Kaplanov I, Carmi Y, Kornetsky R, Shemesh A, Shurin GV, Shurin MR, Dinarello CA, Voronov E, Apte RN. Blocking IL1β reverses the immunosuppression in mouse breast cancer and synergizes with anti-PD-1 for tumor abrogation. Proc. Natl. Acad. Sci. USA 116, 1361–1369 (2019).

19. Zhang Y, Chen X, Wang H, Gordon-Mitchell S, Sahu S, Bhagat TD, Choudhary G, Aluri S, Pradhan K, Sahu P, Carbajal M, Zhang H, Agarwal B, Shastri A, Martell R, Starczynowski D, Steidl U, Maitra A, Verma A. Innate immune mediator, Interleukin-1 receptor accessory protein (IL1RAP), is expressed and pro-tumorigenic in pancreatic cancer. J. Hematol. Oncol. 15, 70 (2022).

20. Lv Q, Xia Q, Li A, Wang Z. The Potential Role of IL1RAP on Tumor Microenvironment-Related Inflammatory Factors in Stomach Adenocarcinoma. Technol. Cancer Res. Treat 20, 5282 (2021).

21. Li F, Zhang W, Wang M, Jia P. IL1RAP regulated by PRPRD promotes gliomas progression via inducing neuronal synapse development and neuron differentiation in vitro. Pathol. Res. Pract. 216, 153141 (2020).

22. Rydberg Millrud C, Deronic A, Grönberg C, Jaensson Gyllenbäck E, von Wachenfeldt K, Forsberg G, Liberg D. Blockade of IL1α and IL1β signaling by the anti-IL1RAP antibody nadunolimab (CAN04) mediates synergistic anti-tumor efficacy with chemotherapy. Cancer Immunol. Immunother. 72, 667–678 (2023).

23. Fields JK, Gyllenbäck EJ, Bogacz M, Obi J, Birkedal GS, Sjöström K, Maravillas K, Grönberg C, Rattik S, Kihn K, Flowers M, Smith AK, Hansen N, Fioretos T, Huyhn C, Liberg D, Deredge D, Sundberg EJ. Antibodies targeting the shared cytokine receptor IL-1 receptor accessory protein invoke distinct mechanisms to block all cytokine signaling. Cell Reports 43(5), 114099 (2024).

24. Grönberg C, Rattik S, Tran-Manh C, Zhou X, Rius Rigau A, Li YN, Györfi AH, Dickel N, Kunz M, Kreuter A, Matei EA, Zhu H, Skoog P, Liberg D, Distler JH, Trinh-Minh T. Combined inhibition of IL-1, IL33 and IL-36 signalling by targeting IL1RAP ameliorates skin and lung fibrosis in preclinical models of systemic sclerosis. Annals of the Rheumatic Diseases 83(9), 1156 – 1168 (2024).

25. Aoki S, Hayakawa M, Ozaki H, Takezako N, Obata H, Ibaraki N, Tsuru T, Tominaga S, Yanagisawa K. ST2 gene expression is proliferation-dependent and its ligand, IL33, induces inflammatory reaction in endothelial cells. Mol. Cell Biochem. 335(1-2), 75–81 (2010).

26. Velcicky J, Cremosnik G, Scheufler C, Meier P, Wirth E, Felber R, Ramage P, Schaefer M, Kaiser C, Lehmann S, Kutil R, Singeisen S, Mueller-Ristig D, Popp S, Cebe R, Lehr P, Kaupmann K, Erbel P, Röhn TA, Giovannoni J, Dumelin CE, Martiny-Baron G. Discovery of selective low molecular weight interleukin-36 receptor antagonists by encoded library technologies. Nat. Commun. 16, 1669 (2025).

27. Cohen RB, Jimeno A, Hreno J, Sun L, Wallén-Öhman M, Millrud CR, Sanfridson A, Garcia-Ribas I. Safety, tolerability, and preliminary efficacy of nadunolimab, an anti-IL-1 receptor accessory protein monoclonal antibody, in combination with pembrolizumab in patients with solid tumors. Invest. New Drugs 43(3), 609–620 (2025).

