## Supplementary material for "Anti-IL1RAP Antibodies for Pan-Inhibition of IL-1 Family Cytokine Signaling in Inflammatory Diseases and Oncology": Figure S1-S6; Table S1-S2

**Supplementary Materials**

**Supplementary Figures**

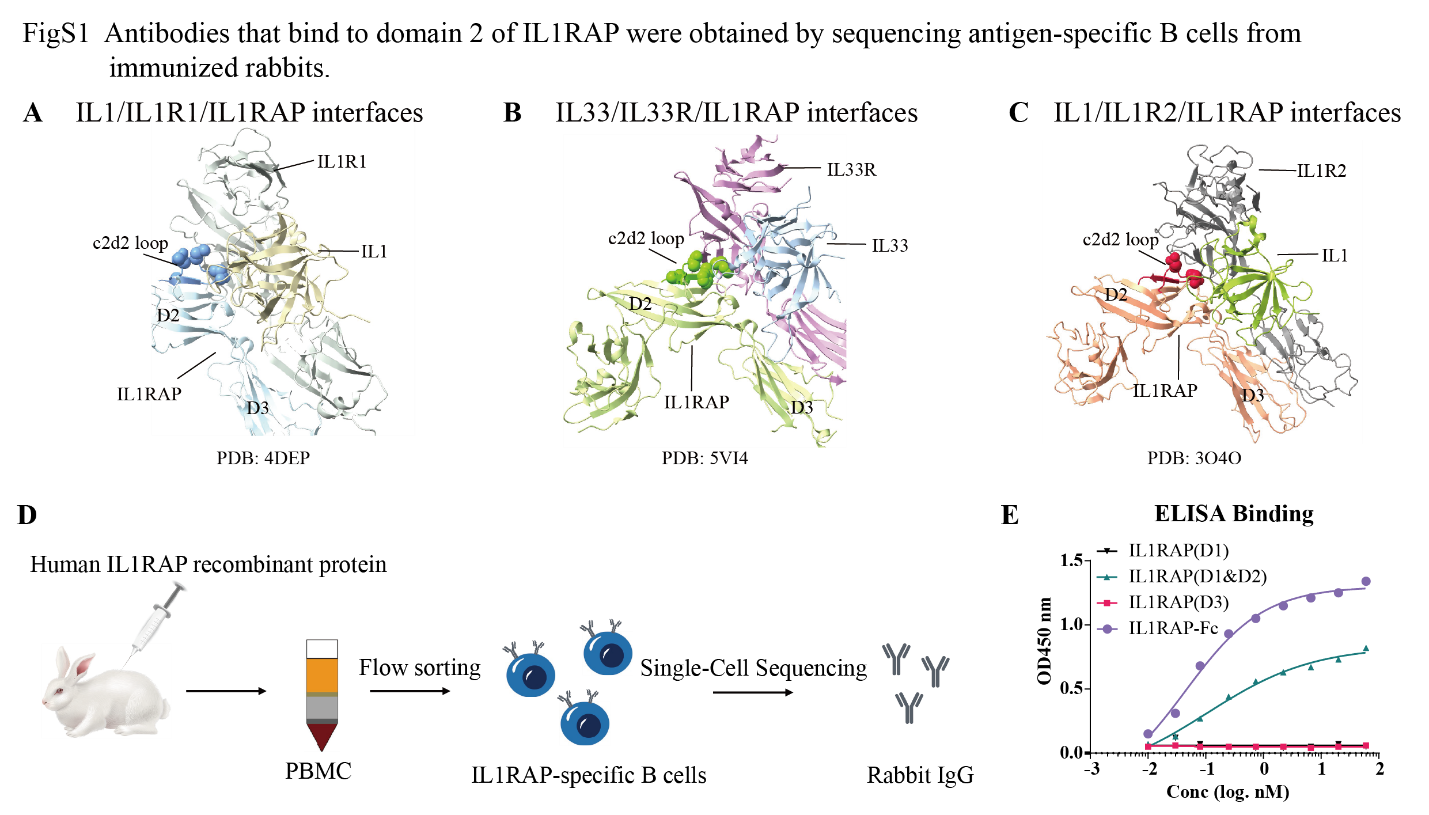
**Figure S1**.

**Fig. S1**. **Antibodies that bind to domain 2 of IL1RAP were obtained by sequencing antigen-specific B cells from immunized rabbits.** (A) Crystal structure of IL1RAP (PDB: 4DEP), with IL1 (olive), IL-1RI (grey) and c2d2 loop (blue) interfaces colored. (B) Crystal structure of IL1RAP (PDB: 5VI4), with IL33 (cyan), IL-33R (pink) and c2d2 loop (green) interfaces colored. (C) Crystal structure of IL1RAP (PDB: 3O4O), with IL1 (dark goldenrod), IL-1R2 (deep pink) and c2d2 loop (red) interfaces colored. (D) Schematic diagram of antibody discovery and screening workflow. (E) ELISA binding assessment of the parental antibody to full-length IL1RAP and its individual domains.

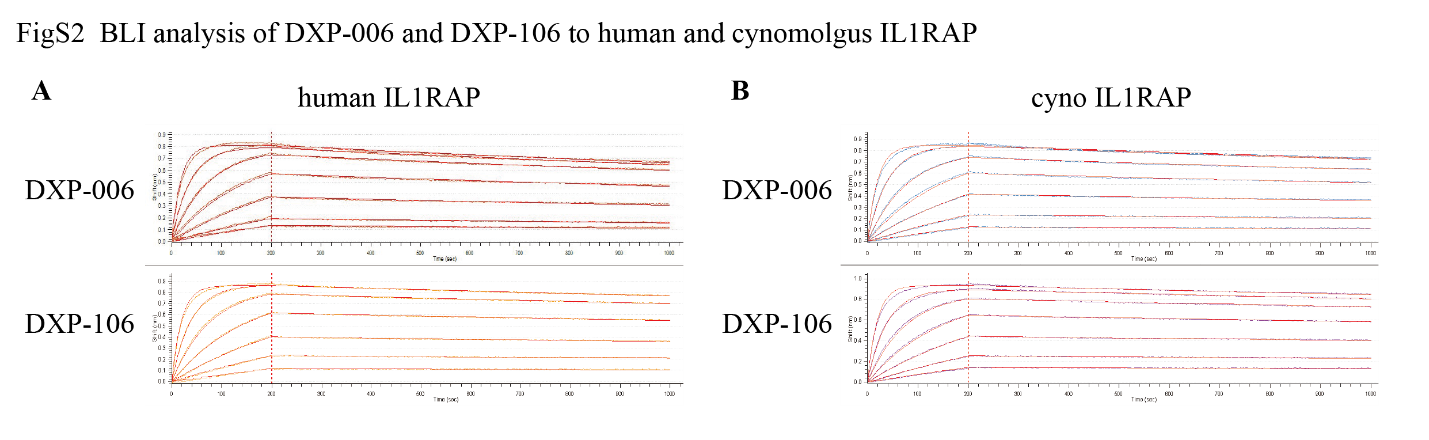
**Figure S2.**

**Fig. S2.** **BLI analysis of DXP-006 and DXP-106 to human and cynomolgus IL1RAP.**

(A) BLI analysis of DXP-006 and DXP-106 to human IL1RAP. (B) BLI analysis of DXP-006 and DXP-106 to cynomolgus IL1RAP.

**Figure S3.**

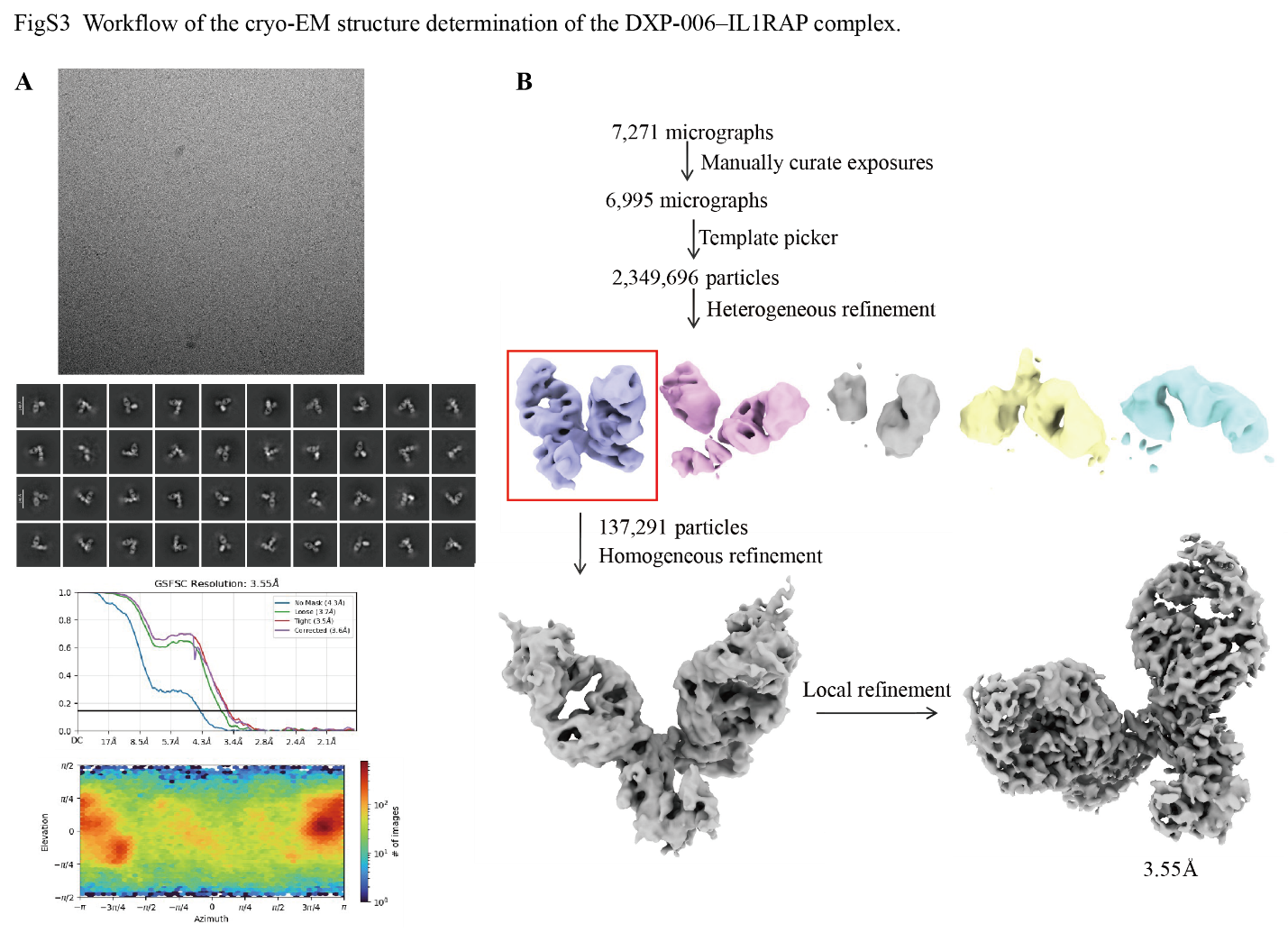
**Fig. S3.** **Workflow of the cryo-EM structure determination of the DXP-006–IL1RAP complex.** (A) Representative cryo-EM micrograph of the DXP-006 Fab–IL1RAP complex acquired on a Titan Krios G4 microscope equipped with a Falcon 4 camera and Selectris X energy filter. (B) 2D class averages showing distinct secondary structural features and particle orientations.

**Figure S4.**

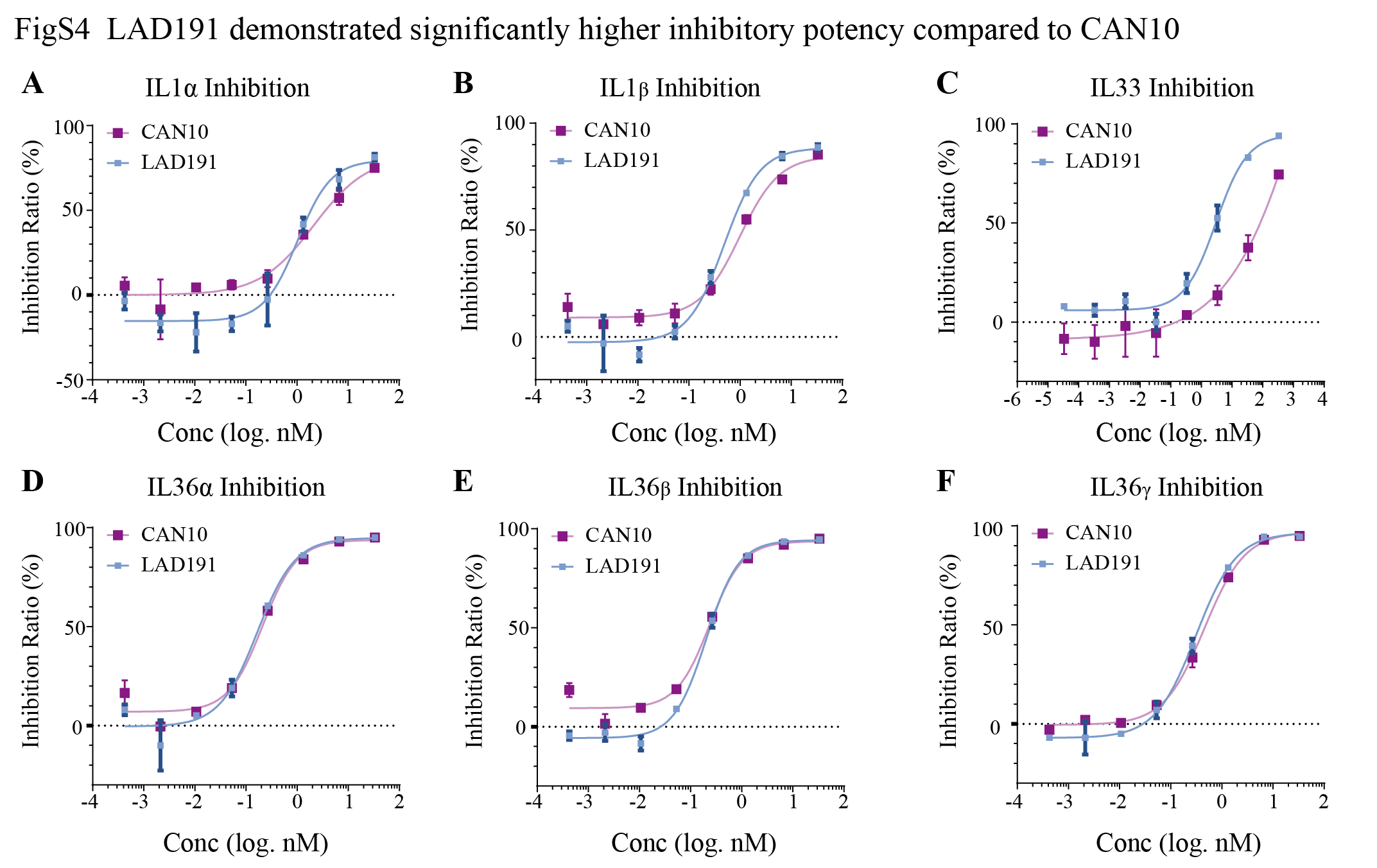
**Fig. S4. LAD191 demonstrated higher inhibitory potency compared to CAN10.** (A-B) Inhibition of IL1α (A) and IL1β (B) induced signaling by indicated antibodies in HEK-Blue IL-1 reporter cells. (C) Inhibition of IL33 induced signaling by indicated antibodies in HUVEC cells. (D-F) Inhibition of IL36α (D), IL36β (E) and IL36γ (F) induced signaling by indicated antibodies in A431 cells.

**Figure S5.**
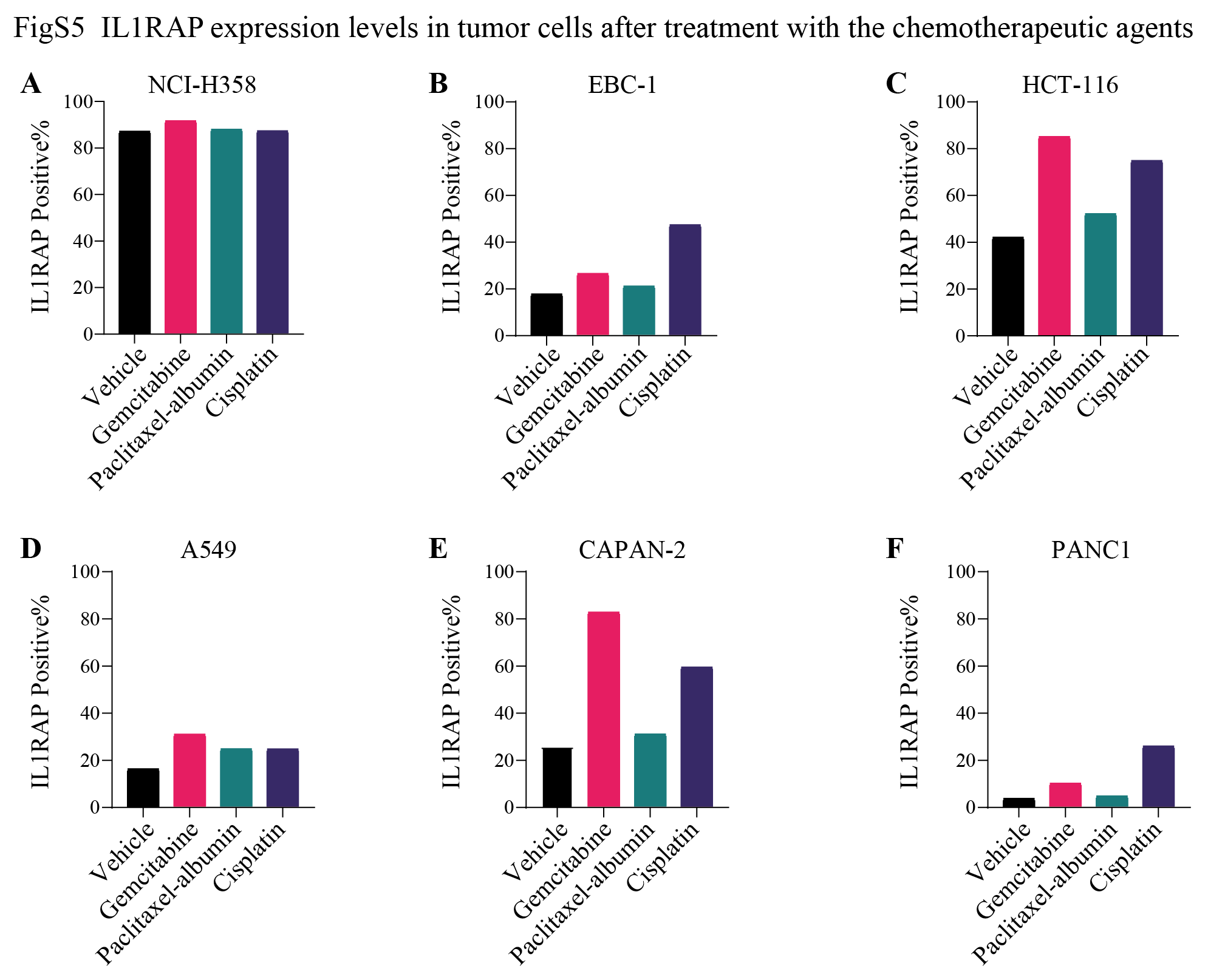

**Fig. S5. IL1RAP levels in tumor cells after treatment with the chemotherapeutic agents.** (A-F) Flow cytometric analysis of IL1RAP surface expression on human cancer cell lines NCI-H358 (A), EBC-1 (B), HCT116 (C), A549 (D), CaPan-2 (3) and PANC1 (F) after 24-hour exposure to gemcitabine, paclitaxel-albumin, or cisplatin.

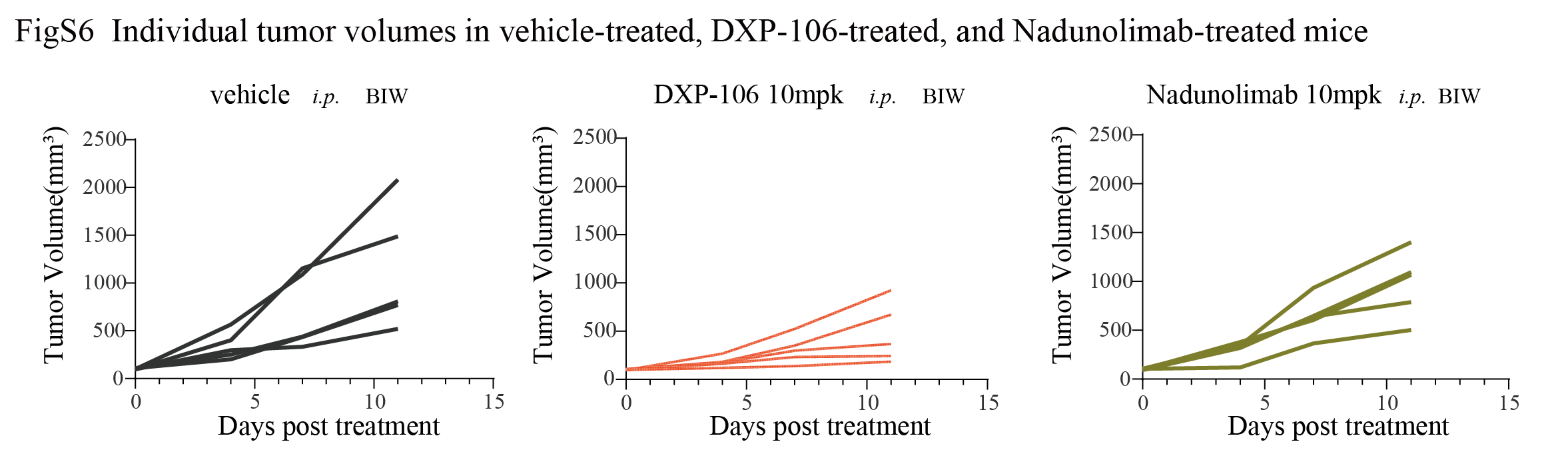
**Figure S6.**

**Fig. S6. Individual tumor volumes in vehicle-treated, DXP-106-treated, and Nadunolimab-treated mice.** Individual tumor growth curves of CB17 SCID mice bearing NCI-H358 subcutaneous xenografts. Mice were administered vehicle (PBS), Nadunolimab (10 mg/kg), or DXP-106 (10 mg/kg) twice weekly via intraperitoneal injection.

**Supplementary Tables**

**Table S1.** **The binding affinity of DXP-006 and DXP-106 to human and cynomolgus IL1RAP.**

| Antibody | Antigen | koff(1/s) | kon(1/Ms) | KD(M) | FullR2 |
| --- | --- | --- | --- | --- | --- |
| DXP-006 | huIL1RAP | 2.57E-04 | 2.73E+05 | 9.42E-10 | 0.9982 |
|  | cynoIL1RAP | 1.91E-04 | 2.51E+05 | 7.61E-10 | 0.9985 |
| DXP-106 | huIL1RAP | 1.46E-04 | 2.58E+05 | 5.67E-10 | 0.9987 |
|  | cynoIL1RAP | 1.27E-04 | 2.62E+05 | 4.86E-10 | 0.9986 |

**Table S2. Cryo-EM data collection, refinement, and validation statistics**

|  | DXP006-IL1RAP complex map |
| --- | --- |
| Data collection and processing  Microscope  Camera  Imaging mode | KriosG4  Falcon 4  Counted resolution |
| Magnification | 130k |
| Voltage (kV) | 300 |
| Electron exposure (e–/Å2) | 60 |
| Defocus range (μm) | -1.0~-2.0 |
| Pixel size (Å) | 0.95 |
| Symmetry imposed | C1 |
| Initial particle images (no.) | 2,349,696 |
| Final particle images (no.) | 137,291 |
| Map resolution (Å)  FSC threshold | 3.55  0.143 |
| Map resolution range (Å) | 2.5-5.0 |
| Refinement |  |
| Initial model used (PDB code) | AF3 predicted |
| Map sharpening B factor (Å2) | -72.4 |
| Model composition  Protein residues  Ligands | 691  1 |
| R.m.s. deviations  Bond lengths (Å)  Bond angles (°)  MolProbity score  Clashscore | 0.004  0.659  2.10  11.52 |
| Validation  Rotamers outliers (%)  Cβ outliers (%) | 0.00  0.00 |
| Ramachandran plot  Favored (%)  Allowed (%)  Disallowed (%) | 91.02  8.98  0.00 |
